# Depletion of bleomycin hydrolase (Blmh) downregulates histone demethylase Phf8, impairs mTOR signaling/autophagy, accelerates amyloid beta accumulation, and induces neurological deficits in mice

**DOI:** 10.1101/2023.03.20.533511

**Authors:** Łukasz Witucki, Kamila Borowczyk, Joanna Suszyńska-Zajczyk, Ewelina Warzych, Piotr Pawlak, Hieronim Jakubowski

## Abstract

Bleomycin hydrolase (BLMH), a homocysteine (Hcy)-thiolactone detoxifying enzyme, is attenuated in brains of Alzheimer’s disease patients. In mice, Blmh depletion causes astrogliosis and behavioral changes. Depletion of histone demethylase PHF8, which controls mTOR signaling by demethylating H4K20me1, causes neuropathy in humans and mice. Here we examined how Blmh depletion affects the Phf8/H4K20me1/mTOR signaling/autophagy pathway and amyloid beta (Aβ) accumulation and cognitive/neuromotor performance in mice. We found that Phf8 was significantly downregulated in brains of *Blmh*^-/-^ mice *vs*. *Blmh*^+/+^ sibling controls. H4K20me1, mTOR, phospho-mTOR, and App were upregulated while autophagy markers Bcln1, Atg5, and Atg7 were downregulated in *Blmh*^-/-^ brains. Blmh depletion caused similar biochemical changes and significantly elevated Aβ in *Blmh*^-/-^5xFAD *vs*. *Blmh*^+/+^5xFAD brains. Behavioral testing identified cognitive/neuromotor deficits in *Blmh*^-/-^ and *Blmh*^-/-^5xFAD mice. In Blmh-depleted N2a-APP_swe_ cells, Phf8 was downregulated, while APP, total H4K20me1, and H4K20me1-*mTOR* promoter binding were elevated. This led to mTOR upregulation, autophagy downregulation, and significantly increased APP and Aβ levels. Phf8 depletion or treatments with Hcy-thiolactone or *N*-Hcy-protein, metabolites that are elevated in Blmh-depleted mice, induced similar biochemical changes in N2a-APP_swe_ cells, akin to those in induced by Blmh depletion. Taken together, our findings indicate that Blmh interacts with APP and the Phf8/H4K20me1/mTOR/autophagy pathway and show that disruption of these interactions lead to Aβ accumulation and cognitive and neuromotor deficits.

## 1 INTRODUCTION

Bleomycin hydrolase (Blmh), named for its ability to deaminate and inactivate the anticancer glycopeptide drug bleomycin, is a thiol-dependent cytoplasmic aminopeptidase expressed in human and rodent organs, including the brain (1, 2). In addition to the aminopeptidase activity, Blmh has a thiolactonase activity and participates in homocysteine (Hcy) metabolism by detoxifying Hcy-thiolactone (3, 4).

Hcy-thiolactone is formed from Hcy in an error-editing reaction in protein biosynthesis catalyzed by methionyl-tRNA synthetase (5). Accumulation of Hcy-thiolactone is harmful because of its ability to modify protein lysine residues (6), which generates structurally- and functionally-impaired *N*-homocysteinylated (*N*-Hcy)-proteins with proinflammatory, prothrombotic, and pro-amyloidogenic properties (7). Hcy-thiolactone and *N*-Hcy-proteins accumulate in mentally retarded CBS- and MTHFR-deficient patients (7) and are mechanistically linked to neurological diseases such as Alzheimer’s disease (AD) (8, 9), stroke (10), cognitive impairment (11), Parkinson’s disease (12), and neural tube defects (13, 14), as well as cardiovascular disease (15), cancer (16–18), and rheumatoid arthritis (19).

BLMH has been linked to AD and Huntington disease (HD). Specifically, BLMH has the ability to process amyloid precursor protein (APP) to amyloid beta (Aβ) (20) and to further process Aβ (21). BLMH has also the ability to generate *N*-terminal fragments of huntingtin, thought to be important mediators of HD pathogenesis (22). In human brain, BLMH is localized in neocortical neurons and in dystrophic neurites of senile plaques (23). A polymorphism in human *BLMH* gene, resulting in I443V substitution in the BLMH protein, is associated with an increased risk of AD in some (24, 25), albeit no other studies (26–28).

The Hcy-thiolactonase and aminopeptidase activities of BLMH are decreased in brains of AD patients, suggesting that the attenuated BLMH activity could contribute to the pathology of AD (29). In mice, deletion of the *Blmh* gene causes astrogliosis and behavioral changes (30). Furthermore, *Blmh*^-/-^ mice exhibit diminished ability to detoxify Hcy-thiolactone, which elevates brain Hcy-thiolactone levels, and increases neurotoxicity of Hcy-thiolactone injections (4). Studies of *Blmh*^-/-^ mouse brain proteome demonstrated that Blmh interacts with diverse cellular processes, such as synaptic plasticity, cytoskeleton dynamics, cell cycle, energy metabolism, and antioxidant defenses that are essential for brain homeostasis (8). Collectively, these findings suggest that Blmh plays an important role in the central nervous system (CNS).

Plant homeodomain finger protein 8 (PHF8) has been identified as one of the X chromosome genes linked to intellectual disability syndrome, autism spectrum disorder, attention deficit hyperactivity disorder (31), and severe mental retardation (32). PHF8 is a histone demethylase that can demethylate H4K20me1, H3K9me2/me1, and H3K27me2. Demethylation of H4K20me1 by PHF8 is important for maintaining homeostasis of mTOR signaling. The phenotype of human PHF8 deficiency has been replicated in *Phf8*^-/-^ mice, which show impaired hippocampal long-term potentiation and behavioral deficits in learning and memory (33).

In the present work we examined a role of Blmh in the CNS by studying consequences of Blmh depletion. Because dysregulated mTOR signaling and autophagy have been implicated in Aβ accumulation in AD brains (34, 35), and H4K20me1 demethylation by PHF8 is important for maintaining homeostasis of mTOR signaling, we examined how these processes are affected in brains of *Blmh*^-/-^ *vs*. *Blmh^+^*^/+^ mice as well as in transgenic *Blmh*^-/-^5xFAD *vs*. *Blmh^+^*^/+^5xFAD mice, which overproduce Aβ. We also examined how changes in these processes and in APP/Aβ expression in Blmh-depleted mice relate to behavioral performance of these mice. We studied underlying molecular mechanisms by manipulating Blmh and Phf8 expression as well as Hcy-thiolactone and *N*-Hcy-protein levels in Aβ-overproducing mouse neuroblastoma N2a-APPswe cells.

## 2 MATERIALS AND METHODS

### 2.1 Mice

*Blmh*^-/-^ (36) and 5xFAD (37) mice on the C57BL/6J genetic background were housed and bred at the Rutgers-New Jersey Medical School Animal Facility. 5xFAD mice overexpress the K670N/M671L (Swedish), I716V (Florida), and V717I (London) mutations in human APP(695), and M146L and L286V mutations in human PS1. 5xFAD mice accumulate high levels of Aβ42 beginning around 2 months of age (37) (https://www.alzforum.org/research-models/5xfad-b6sjl). The *Blmh*^-/-^ mice were crossed with 5xFAD animals to generate *Blmh*^-/-^5xFAD mice and their *Blmh^+^*^/+^5xFAD sibling controls. Mouse *Blmh* genotype were established by PCR of tail clips using the following primers: *Blmh* intron 2 forward primer p1 (5′-CACTGTAGCTGTACTCACAC), *Blmh* exon 3 reverse primer p2 (5′-GCGACAGAGTACCATGTAGG-3′) and neomycin cassette reverse primer p3 (5′-ATTTGTCACGTCCTGCACGACG-3′) (36). 5xFAD genotype was established using human APP and PS1 primers (hAPP forward 5′-AGA GTA CCA ACT TGC ATG ACT ACG-3′ and reverse 5′-ATG CTG GAT AAC TGC CTT CTT ATC-3′; hPS1 forward 5′-GCT TTT TCC AGC TCT CAT TTA CTC-3′ and reverse 5′-AAA ATT GAT GGA ATG CTA ATT GGT-3′). The mice were fed with a standard rodent chow (LabDiet5010; Purina Mills International, St. Louis MO, USA) (4). Two- and four-month-old *Blmh*^-/-^ mice and their *Blmh*^+/+^ siblings, as well as 5- and 12-month-old *Blmh*^-/-^5xFAD mice and their *Blmh^+^*^/+^5xFAD siblings were used in experiments. Water supplemented with 1% methionine (a high Met diet) was provided to mice starting at 1 month of age to induce hyperhomocysteinemia, as needed. The high Met diet significantly increases plasma total Hcy levels (*P* < 1.E-06) (6-fold from 6.8 to 39 μM in *Blmh*^-/-^ mice and 10-fold from 7.4 to 77 μM in *Blmh*^+/+^ mice) as well as *N*-Hcy-protein levels (*P* < 0.001) (3-fold from 2.8 to 8.4 μM in *Blmh*^-/-^ mice and 4.5-fold from 1.2 to 5.4 μM in *Blmh*^+/+^ mice) (4). Animal procedures were approved by the Institutional Animal Care and Use Committee at Rutgers-New Jersey Medical School.

### 2.2 Brain protein extraction

Mice were euthanized by CO_2_ inhalation, the brains collected and frozen on dry ice. Frozen brains were pulverized with dry ice using a mortar and pestle and stored at −80°C. Proteins were extracted from the pulverized brains (50±5 mg; 30±3 mg brain was used for Aβ analyses) using RIPA buffer (4 v/w, containing protease and phosphatase inhibitors) with sonication (Bandelin SONOPLUS HD 2070) on wet ice (three sets of five 1-s strokes with 1 min cooling interval between strokes). Brain extracts were clarified by centrifugation (15,000 g, 30 min, 4°C) and clear supernatants containing 8-12 mg protein/mL were collected (RIPA-soluble fraction). Protein concentrations were measured with BCA kit (Thermo Scientific).

For Aβ analyses, pellets remaining after protein extraction with RIPA buffer were re-extracted by brief sonication in 2% SDS, centrifuged (15,000 g, 15 min, room temperature), and the supernatants again collected (SDS-soluble fraction). The SDS-extracted pellets were then extracted by sonication in 70% formic acid (FA), centrifuged, and the supernatants were collected (the FA-soluble fraction) (38).

### 2.3 Cell culture and treatments

Mouse neuroblastoma N2a-APPswe cells, harboring a human APP transgene with the K670N and M671L Swedish mutations (39), and N2a cells without the transgene were grown (37°C, 5% CO_2_) in DMEM/F12 medium (Thermo Scientific) supplemented with 5% FBS, non-essential amino acids, and antibiotics (penicillin/streptomycin) (MilliporeSigma).

After cells reached 70-80% confluency, the monolayers were washed twice with PBS and overlaid with DMEM medium without methionine (Thermo Scientific), supplemented with 5% dialyzed fetal bovine serum (FBS) (MilliporeSigma) and non-essential amino acids. L-Hcy-thiolactone (MilliporeSigma) or *N*-Hcy-protein, prepared as described in ref. (40), were added (at concentrations indicated in figure legends) and the cultures were incubated at 37°C in 5% CO_2_ atmosphere for 24 h.

For gene silencing, siRNAs targeting the *Blmh* (Cat. # 100821 and s63474) or *Phf8* gene (Cat. # S115808, and S115809) (Thermo Scientific) were transfected into cells maintained in Opti-MEM medium by 24-h treatments with Lipofectamine RNAiMax (Thermo Scientific). Cellular RNA for RT-qPCR analyses were isolated as described in section 2.5 below. For protein extraction, RIPA buffer (MilliporeSigma) was used according to manufacturer’s protocol.

### 2.4 Western blots

Proteins were separated by SDS-PAGE on 10% gels (20 µg protein/lane) and transferred to PVDF membrane (Bio-Rad) for 20 min at 0.1 A, 25 V using Trans Blot Turbo Transfer System (Bio-Rad). After blocking with 5 % bovine serum albumin in TBST buffer (1 h, room temperature), the membranes were incubated overnight at 4°C with anti-Blmh (Abcam, AB188371), anti-Phf8 (Abcam, ab36068), anti-H4K20me1 (Abcam ab177188), anti-mTOR (CS #2983), anti-pmTOR Ser2448 (CS, #5536), anti-Atg5 (CS, #12994), anti-Atg7 (CS, #8558), anti-Beclin-1 (CS, #3495), anti-p62 (CS, #23214), anti-Gapdh (CS, #5174), or anti-App (Abcam, ab126732) for 1 hour. Membranes were washed three times with 1X Tris-Buffered Saline, 0.1% Tween 20 Detergent (TBST), 10 min each, and incubated with goat anti-rabbit IgG secondary antibody conjugated with horseradish peroxidase. Positive signals were detected using Western Bright Quantum-Advansta K12042-D20 and GeneGnome XRQ NPC chemiluminescence detection system. Bands intensity was calculated using Gene Tools program from Syngene.

For Western blots analyses of Aβ, brain protein extracts (2 μL) were separated on 10 % Tricine gels, and then transferred (0.5 A, 25 V 10 min) onto 22 µm PVDF membranes (Bio-Rad). The membranes were washed 3 times with 1x TBST and then blocked with 5 % bovine serum albumin (BSA) for 1 h at RT. After blocking, membranes were washed 3 times with 1x TBST and then incubated with primary anti-Aβ antibody (D54D2, CS #8243). Membranes were washed 3 times with 1x TBS-T and incubated with anti-rabbit IgG HRP-linked antibodies (CS#7074) for 1 h at RT. Signals were collected using clarity Max Western ECL Substrate (Bio-Rad) and GeneGnome XRQ - Chemiluminescence imaging (Syngene).

### 2.5 RNA isolation, cDNA synthesis, RT-qPCR analysis

Total RNA was isolated using Trizol reagent (MilliporeSigma). cDNA synthesis was conducted using Revert Aid First cDNA Synthesis Kit (Thermo Fisher Scientific) according to manufacturer’s protocol. Nucleic acid concentration was measured using NanoDrop (Thermo Fisher Scientific). RT-qPCR was performed with SYBR Green Mix and CFX96 thermocycler (Bio-Rad). The 2^(-ΔΔCt)^ method was used to calculate the relative expression levels (41). Data analysis was performed with the CFX Manager™ Software, Microsoft Excel, and Statistica. RT-qPCR primer sequences are listed in **Table S1**.

### 2.6 Chromatin immunoprecipitation assay

For CHIP assays we used CUT&RUN Assay Kit #86652 (Cell Signaling Technology, Danvers, MA, USA) following the manufacturer’s protocol. Each ChIP assay was repeated 3 times. Briefly, for each reaction we used 100 000 cells. Cells were trypsinized and harvested, washed 3x in ice-cold PBS, bound to concanavalin A-coated magnetic beads for 5 min, RT. Cells were then incubated (4h, 4^◦^C) with 2.5 µg of anti-PHF8 antibody (Abcam, ab36068) or anti-H4K20me1 antibody (Abcam, ab177188) in the antibody-binding buffer plus digitonin that permeabilizes cells. Next, cells are treated with pAG-MNase (1 h, 4^◦^C), washed, and treated with CaCl2 to activate DNA digestion (0.5 h, 4^◦^C). Cells were then treated with the stop buffer and spike-in DNA was added for each reaction for signal normalization, and incubated (30 min, 37^◦^C). Released DNA fragments were purified using DNA Purification Buffers and Spin Columns (CS #14209) and quantified by RT-qPCR using primers targeting the promoter, upstream, and downstream regions of the *mTOR* gene (**Table S1**).

### 2.7 Confocal microscopy, Aβ staining in N2a-APPswe cells

Mouse neuroblastoma N2a-APPswe cells were cultured in Millicell EZ SLIDE 8-well glass slides (Merck). After treatments cells were washed with PBS (3 times, 10 minutes each). Cells were fixed with 4% PFA (Sigma-Aldrich) (37^◦^C, 15 min). After fixation, cells were again washed 3 times with PBS buffer and permeabilized in 0.1% Triton X-100 solution (RT, 20 min), blocked with 0.1% BSA (RT, 1h), and incubated with anti-Aβ antibody (CS #8243; 4^◦^C, 16 h). Cells were then washed 3 times with PBS and stained with secondary antibody Goat Anti-Rabbit IgG H&L (Alexa Fluor® 488) (Abcam, ab150077; RT, 1 h) to detect Aβ. DAPI (Vector Laboratories) was used to visualize nuclei. Fluorescence signals were detected by using a Zeiss LSM 880 confocal microscope with a 488 nm filter for the Alexa Fluor® 488 (Aβ) and 420–480 nm filter for DAPI, taking a *z* stack of 20-30 sections with an interval of 0.66 μm and a range of 15 μm. Zeiss Plan-Apochromat X40/1.2 Oil differential interference contrast objective were used for imaging. Images were quantified with the ImageJ Fiji software (NIH).

### 2.8 Behavioral testing

#### Novel Object Recognition test

NOR is a test of recognition memory (42). The test was carried out in two sessions, divided by a 6-h intersession interval. During the first session (familiarization session), the animal was free to explore two similar objects, and during the second session (test session), one of the objects was replaced by a novel, unfamiliar object. No habituation phase was performed. A minimal exploration time for both objects during both the familiarization and test phase (∼20 s) was used, with a maximal time of 10 min to reach the criterion. Mice were tested in a white plastic box (33 × 33 × 20 cm). W used objects that differ in shape and texture: towers of Lego bricks (8-cm high and 3.2-cm wide, built-in blue, yellow, red, and green bricks) and Falcon tissue culture flasks filled with sand (9.5 cm high, 2.5 cm deep and 5.5 cm wide, transparent plastic with a yellow bottle cap). We scored object exploration whenever the mouse sniffed the object or touched the object while looking at it (i.e., when the distance between the nose and the object was less than 2 cm). Climbing onto the object (unless the mouse sniffs the object it has climbed on) or chewing the object did not qualify as exploration.

#### Hindlimb test

The hindlimb clasping test is used to assess neurodegeneration in mouse models (43). For this test, mice were suspended by the base of the tail and videotaped for 10 seconds. Three separate trials were taken over three consecutive days. Hindlimb clasping was scored from 0 to 3: 0 = hindlimbs splayed outward and away from the abdomen, 1 = one hindlimb retracted inwards towards the abdomen for at least 50% of the observation period, 2 = both hindlimbs partially retracted inwards towards the abdomen for at least 50% of the observation period, 3 = both hindlimbs completely retracted inwards towards the abdomen for at least 50% of the observation period. Hindlimb clasping scores were added together for the three separate trials.

#### Ledge test

The ledge test is used to assess motor deficits in rodent models of CNS disorders (44). Typically, mice walk along the ledge of a cage and try to descend back into the cage. Three separate trials were taken for each mouse. Ledge test was scored from 0 to 3 points: 0 = a mouse walked along the ledge without slipping and lowered itself back into the cage using paws; 1 = the mouse lost its footing during walking along the ledge but otherwise appeared coordinated; 2 = the mouse did not effectively use its hind legs and landed on its head rather than paws when descending into the cage; 3 = the mouse fell of the ledge or was shaking, barely moving.

#### Cylinder test

The cylinder test is used to assess sensorimotor function in rodent models of CNS disorders. A mouse was place in the transparent 500 ml plastic cylinder. The number of times the mouse rears up and touches the cylinder wall during a period of 3 min was counted. A rear is defined as a vertical movement with both forelimbs off the floor so that the mouse is standing only on its hindlimbs. At the end of 3 min, the mouse was removed and placed back into its home cage. Because spontaneous activity in the cylinder is affected by repeated testing resulting in reduced activity over time, mice were only once in their lifetime.

### 2.9 Statistical analysis

The results were calculated as mean ± standard deviation. A two-sided unpaired t test was used for comparisons between two groups of variables, *P* < 0.05 was considered significant. Statistical analysis was performed using Statistica, Version 13 (TIBCO Software Inc., Palo Alto, CA, USA, http://statistica.io).

## 3 RESULTS

### 3.1 Blmh depletion downregulates histone demethylase Phf8 and upregulates H4K20me1 epigenetic mark in the mouse brain

To determine if Blmh interacts with Phf8, we quantified Phf8 protein in brains of young *Blmh*^-/-^ mice and their *Blmh*^+/+^ sibling controls by Western blotting. We also examined effects of hyperhomocysteinemia (HHcy), induced by providing 1% methionine in drinking water, on the Blmh-Phf8 interaction. We found that Phf8 protein was significantly downregulated in brains of *Blmh*^-/-^ mice *vs*. *Blmh*^+/+^ sibling controls in mice fed with a standard chow diet (from 1.0±0.1 to 0.58±0.10 and 0.62±0.07 in 2- and 4-months-old mice, *P*_genotype_ = 4.E-09 and 4.E-07, respectively; **Figure 1A**). Reduced expression of Phf8 in *Blmh*^-/-^ *vs*. *Blmh*^+/+^ brains was also observed in mice fed with HHcy diet (from 0.80±0.06 to 0.63±0.10 and from 0.67±0.11 to 0.55±0.07 in 2- and 4-months-old mice, *P*_genotype_ = 0.001 and 0.013, respectively; **Figure 1A**).

**Figure 1.**
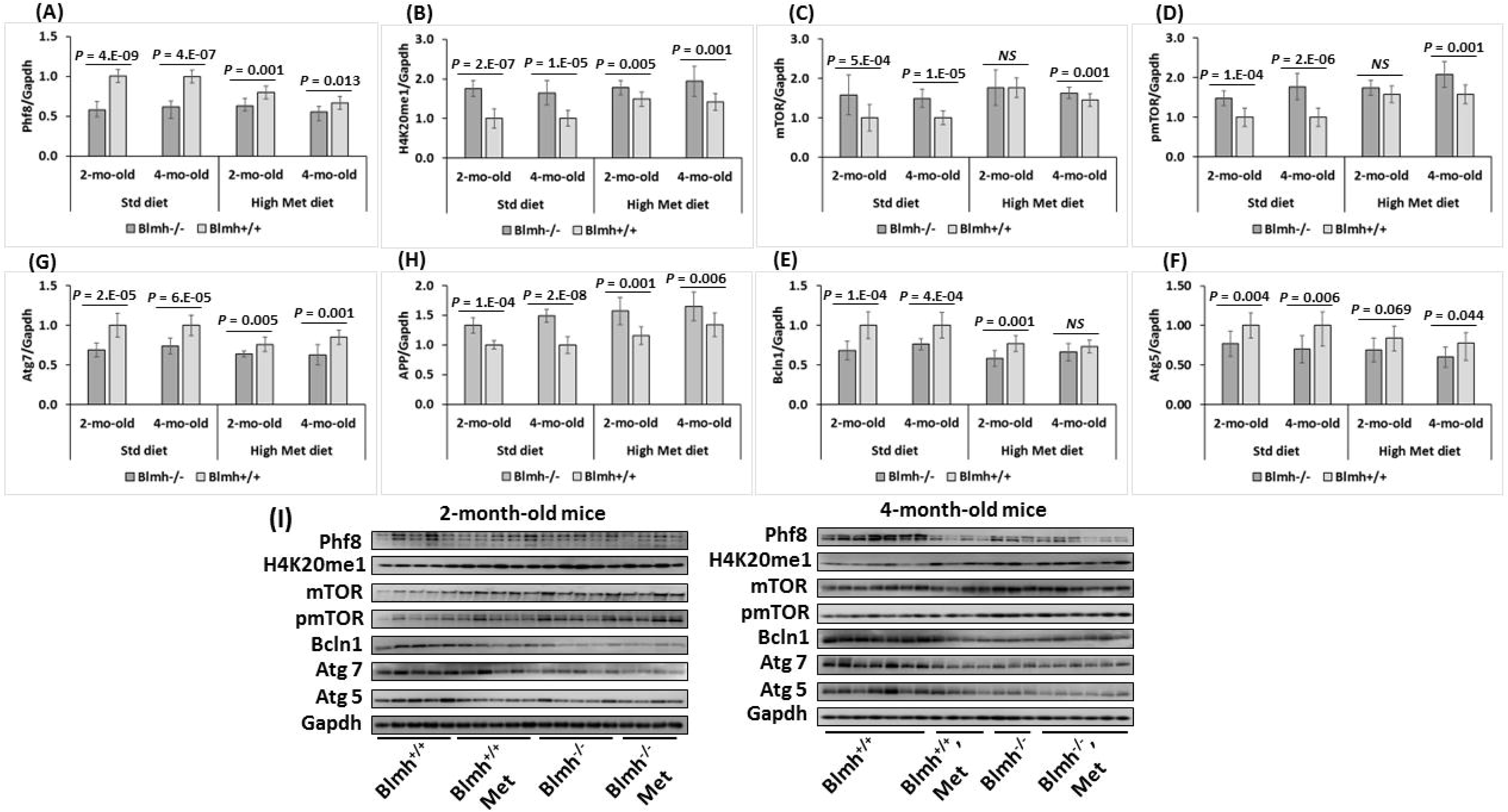
Blmh depletion affects the expression of histone demethylase Phf8, histone H4K20me1 epigenetic mark, mTOR signaling, autophagy, and App in the *Blmh*^-/-^ mouse brain. One-month-old *Blmh*^-/-^ mice (n = 7 and 14) and *Blmh*^+/+^ sibling controls (n = 10 and 10) fed with HHcy or control diet for one or three months were used in experiments. Dietary HHcy was induced by providing 1% methionine in drinking water. Each group included 7-14 mice of both sexes. Bar graphs illustrating quantification of the following brain proteins by Western blotting are shown: Phf8 (**A**), H4K20me1 (**B**), mTOR (**C**), pmTOR (**D**), Bcln1 (**E**), Atg5 (**F**), Atg7 (**G**), and App (**H**). Gapdh protein were used as references for normalization. Pictures of Western blots used for protein quantification are shown in panel (**I**). Data are averages of three independent experiments.

However, Phf8 expression in *Blmh*^-/-^ mice was essentially not affected by HHcy diet (*P*_diet_ = 0.056 – 0.210) (**Figure 1A**). In contrast, HHcy diet significantly downregulated Phf8 levels in *Blmh*^+/+^ mice (from 1.0±0.1 to 0.63±0.06 and 0.67±0.11 in 2- and 4-months-old mice, *P*_diet_ = 1.E-04 and 1.E-05, respectively).

The histone H4K20me1 epigenetic mark was significantly upregulated in *Blmh*^-/-^ *vs*. *Blmh*^+/+^ brains in mice fed with a standard chow diet (1.7-fold in 2- and 4-months-old mice, *P*_genotype_ = 2.E-07 and 1.E-05, respectively; **Figure 1B**) or HHcy diet (1.2- and 1.4- fold in 2- and 4-months-old mice, *P*_genotype_ = 0.005 and 0.001, **Figure 1B**). HHcy diet significantly elevated H4K20me1 levels in 2- and 4-months-old *Blmh*^+/+^ mice (1.5- and 1.4-fold, *P*_diet_ = 2.E-07 and 0.005, respectively) and but not in *Blmh*^-/-^ animals (*P*_diet_ = 0.080 – 0.873) **(Figure 1B)**.

To determine how the Blmh-Phf8 interaction is affected by Aβ accumulation, we quantified Phf8 in brains of young (5-month-old) and old (12-month-old) Aβ-overproducing *Blmh*^-/-^5xFAD mice. We found significant effects of Blmh depletion on Phf8 (**Figure S1A**) and H4K20me1 (**Figure S1A**) in 5- and 12-month-old Aβ-overproducing *Blmh*^-/-^5xFAD mice, like those observed in young *Blmh*^-/-^ animals (**Figure 1A, B**). HHcy diet abrogated effects of Blmh depletion on Phf8 and H4K20me1 in 12-month-old but not in 5-month-old *Blmh*^-/-^5xFAD mice (**Figure S1A, B**).

### 3.2 Blmh depletion upregulates mTOR signaling and inhibits autophagy in mouse brain

Because Phf8/H4K20me1 regulate mTOR signaling, we next examined effects of Blmh depletion on levels of mTOR and its active form, phosphorylated at Ser2448 (pmTOR). We found that mTOR protein was significantly upregulated in brains of *Blmh*^-/-^ *vs*. *Blmh*^+/+^ mice (1.6- and 1.5-fold in 2- and 4-months-old, *P*_genotype_ = 0.006 and 1.E-05, respectively; **Figure 1C**). Effect of *Blmh*^-/-^ genotype on mTOR expression was abrogated by HHcy diet (*P*_genotype_ = 0.47) or attenuated (to 1.1-fold, *P*_genotype_ = 5.E-04) in 2-month- or 4-month-old mice, respectively.

HHcy diet significantly increased mTOR expression in 2-month-old *Blmh*^-/-^ (*P*_diet_ =0.008) and in 2- and 4-month-old *Blmh*^+/+^ mice (1.8- and 1.5-fold in 2- and 4-months-old mice, respectively; *P*_diet_ < 5.E-05). However, HHcy diet did not affect mTOR levels in 4-month-old animals (*P*_diet_ = 0.128) (**Figure 1C**).

As mTOR signaling is activated by phosphorylation, we quantified mTOR phosphorylated at Ser2448 (pmTOR). We found that pmTOR was significantly elevated in brains of *Blmh*^-/-^ *vs*. *Blmh*^+/+^ mice (1.5 – 1.8-fold in 2- and 4-month-old mice, *P*_genotype_ = 1.E-04 and 2.E-06, respectively *P*; **Figure 1D**). Effect of *Blmh*^-/-^ genotype on pmTOR levels was abrogated by HHcy diet (*P*_genotype_ = NS) or attenuated (*P*_genotype_ = 0.001) in 2- or 4-month-old mice, respectively.

HHcy diet elevated pmTOR levels in 2- and 4-month-old *Blmh*^-/-^ mice (2-month-old: from 1.5 to 1.7, *P_diet_* = 0.008; 4-month-old: from 1.8 to 2.1, *P_diet_* = 0.049) and *Blmh*^+/+^ mice (2- and 4-month-old: from 1.0 to 1.6, *P_diet_* = 1.E-05) (**Figure 1D**). These findings indicate that Blmh depletion upregulated pmTOR to a similar extent as mTOR.

Because mTOR is a major regulator of autophagy, we quantified effects of Blmh depletion on autophagy-related proteins. We found that the regulators of autophagosome assembly Bcln1, Atg5, and Atg7 were significantly downregulated in brains of *Blmh*^-/-^ *vs*. *Blmh*^+/+^ mice (by 24-39%, *P*_genotype_ = 2.E-05 to 0.006, **Figure 1E, F, G**). Effects of *Blmh*^-/-^ genotype on autophagy were attenuated (Atg7) or abrogated (Beclin1, Atg5) by HHcy diet.

HHcy diet significantly downregulated Bcln1, Atg5, and Atg7 levels in *Blmh*^+/+^ mice (by 45-76%, *P_diet_* = 0.001 – 0.045) but not in *Blmh*^-/-^ animals (*P_diet_* = NS) (**Figure 1E, F, G**). These findings indicate that *Blmh*^-/-^ genotype impaired autophagy regardless of diet. However, HHcy diet impaired autophagy only in *Blmh*^+/+^ mice.

We found that Blmh depletion similarly upregulated mTOR and pmTOR levels in young (5-month-old) Aβ-overproducing *Blmh*^-/-^5xFAD mice (**Figure S1C, D**), like in *Blmh*^-/-^ mice (**Figure 1C, D**). These effects of Blmh depletion disappeared in old (12-month) *Blmh*^-/-^5xFAD mice (**Figure S1C, D**).

Blmh depletion downregulated autophagy-related proteins Beclin1, Atg5, and Atg7 in young and old (5- and 12-month-old, respectively) Aβ-overproducing *Blmh*^-/-^5xFAD mice (**Figure S1E, F, G**), like in young *Blmh*^-/-^ mice (**Figure 1E, F, G**). We also found that p62, a receptor for degradation of ubiquitinated substrates, was upregulated in 5-month-old, and to a lesser extent in 12-month-old, *Blmh*^-/-^5xFAD mice (**Figure S1I**). HHcy diet attenuated or abrogated effects of Blmh depletion on p62 (**Figure S1I**) or Atg7 (**Figure S1G**), but not on Bcln1 (**Figure S1E**) and Atg5 (**Figure S1F**).

### 3.3 Blmh depletion upregulates App expression in mouse brain

We found that App was significantly elevated in 2-month-old *Blmh*^-/-^ *vs*. *Blmh*^+/+^ mice (1.3-fold, *P_genotype_* = 1.E-06). We found a similar elevation in App in 4-month-old *Blmh*^-/-^ *vs*. *Blmh*^+/+^ mice (1.5-fold, *P_genotpe_* = 2.E-08) (**Figure 1H**). App was similarly upregulated by Blmh depletion in mice fed with HHcy diet (1.35- and 1.23-fold in 2- and 4-month-old mice, *P_genotpe_* = 0.001 and 0.006, respectively. HHcy diet upregulated App in *Blmh*^-/-^ mice (1.6-fold in 2- and 4-month-old, *P_diet_* = 0.013 – 0.093) and to a lesser extent in *Blmh*^+/+^ animals (1.13-fold in 2-month-old, *P_diet_* = 0.006 and 1.34-fold in 4-month-old, *P_diet_*= 1.E-04) (**Figure 1H**). We observed similar effects of Blmh depletion on App in Aβ-overproducing 12-month-old *Blmh*^-/-^5xFAD mice (2.0-fold, *P_genotype_* = 1.E-05), which were abrogated by HHcy diet (*P_genotype_* = 0.130) (**Figure S1H**).

### 3.4 Blmh depletion by RNA interference downregulates histone demethylase Phf8, upregulates H4K20me1 epigenetic mark, mTOR signaling, APP, and inhibits autophagy in mouse neuroblastoma N2a-APPswe cells

To elucidate the mechanism by which Blmh depletion impacts Phf8 and its downstream effects on mTOR, autophagy, and APP we first examined whether the findings in *Blmh*^-/-^ mice can be recapitulated in cultured mouse neuroblastoma N2a-APPswe cells that overexpress mutated human APP. We silenced the *Blmh* gene in N2a-APPswe cells by RNA interference using *Blmh*-targeting siRNA and studied how the silencing impacts Phf8 and its downstream effects. Changes in examined protein and mRNA levels in *Blmh*-silenced and control cells were analyzed by Western blotting and RT-qPCR using Gapdh protein and mRNA, respectively, as a reference.

We found that the Blmh protein level was reduced by 80% in *Blmh*-silenced cells (*P* = 4.E-06; **Figure S2A**). We also found that the histone demethylase Phf8 protein level was also significantly reduced (by 40%, *P* = 1.E-06; **Figure S2B**), while the histone H4K20me1 level was significantly elevated (∼2-fold, *P* = 4.E-06; **Figure S2C**) in *Blmh*-silenced N2a-APPswe cells.

At the same time, mTOR protein was significantly upregulated in *Blmh*-silenced N2a-APPswe cells (∼1.7-fold, *P* = 4.E-07; **Figure S2D)**, as was pmTOR (∼1.8-fold, *P* = 3.E-08; **Figure S2E**) and APP (1.6-fold, *P* = 1.E-05; **Figure S2I**), while autophagy-related proteins Bcln1, Atg5, and Atg7 (**Figure S2F, S2G,** and **S2H**, respectively) were significantly downregulated (by 27-36%, *P* = 7.E-08 to 5.E-06).

We found similar changes in the levels of corresponding mRNAs in *Blmh*-silenced N2a-APPswe cells (**Figure S3**). Specifically, Blmh mRNA level was significantly reduced (by 95%, *P* = 0.001; **Figure S3A**), as were Phf8 mRNA (by 35-40%, *P* = 0.005; **Figure S3B**) and mRNAs for autophagy-related proteins Bcln1, Atg5, and Atg7 (by 25-40%, *P* = 0.001 – 0.043; **Figure S3D**, **S3E**, and **S3F**, respectively). mTOR mRNA was significantly upregulated (2.0-2.2-fold, *P* = 0.019; **Figure S3C**) as was APP mRNA (1.3-1.7-fold, *P* = 0.004 – 0.035; **Figure S3G**) in *Blmh*-silenced N2a-APPswe cells, reflecting changes in the corresponding protein levels in these cells (**Figure S2A-I**). These findings indicate that *Blmh* gene exerts transcriptional control over the expression of Phf8, mTOR, APP, Bcln1, Atg5, and Atg7.

The Western blot (**Figure S2**) and RT-qPCR (**Figure S3**) results show that the changes in Phf8, H4K20m31, mTOR signaling, autophagy, and APP induced by *Blmh* gene silencing in N2a-APPswe cells recapitulate the *in vivo* findings in the *Blmh*^-/-^ mouse brain (**Figure 1**).

### 3.5 Hcy-thiolactone and *N*-Hcy-protein downregulate Phf8 histone demethylase, upregulate APP, the H4K20me1 epigenetic mark, and mTOR, and impair autophagy in N2a-APPswe cells

Effects of Blmh depletion on Phf8 and its downstream targets can be caused by the absence the Blmh protein or by upregulation of Hcy-thiolactone and *N*-Hcy-protein, which are known to be elevated in *Blmh*^-/-^ mice (4). To distinguish between these possibilities, we treated N2a-APPswe cells with different concentrations of Hcy-thiolactone or *N*-Hcy-protein.

We found significantly reduced Phf8 levels in N2a-APPswe cells treated with 20 μM Hcy-thiolactone (0.66±0.05) or 10 μM *N*-Hcy-protein (0.77±0.04) compared to untreated control cells (1.00±0.09, *P* < 0.005; **Figure 2A**). Levels of the histone H4K20me1 mark were significantly elevated in N2a-APPswe cells treated with 20 μM Hcy-thiolactone (1.48±0.31) or 10 μM *N*-Hcy-protein (1.42±0.11) compared to untreated control cells (1.00±0.17, *P* < 0.001; **Figure 2B**).

**Figure 2.**
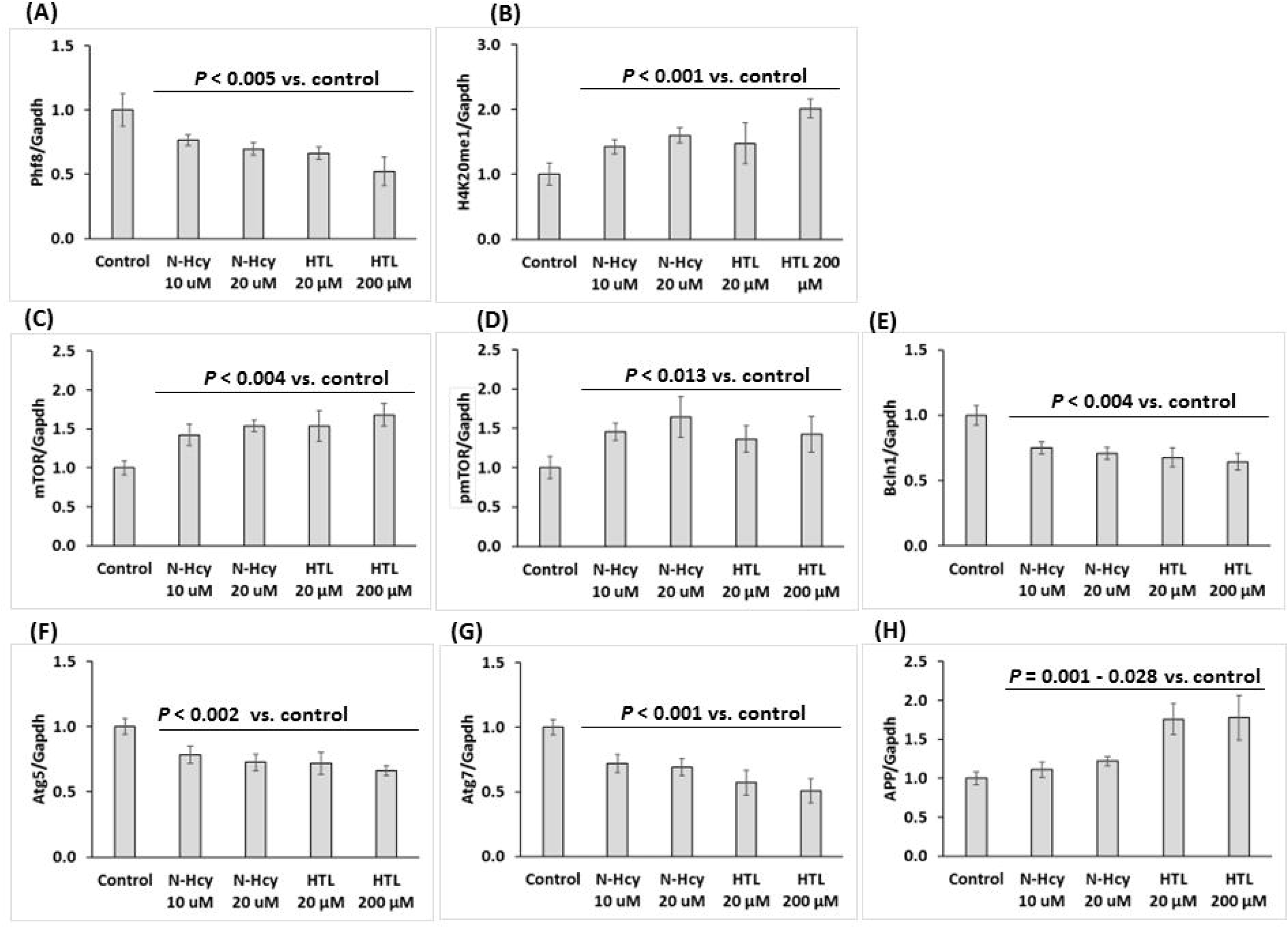
Hcy-thiolactone and *N*-Hcy-protein downregulate Phf8, upregulate H4K20me1 epigenetic mark, mTOR signaling, APP, and impair autophagy in mouse neuroblastoma N2a-APPswe cells. N2a-APPswe cells were treated with indicated concentrations of *N*-Hcy-protein (N-Hcy) or Hcy-thiolactone (HTL) for 24 h at 37^◦^C. Bar graphs illustrating the quantification of Phf8 (**A**), H4K20me1 (**B**), mTOR (**C**), pmTOR (**D**), Bcln1 (**E**), Atg5 (**F**), Atg7 (**G**), and App (**H**) based on Western blot analyses are shown. Gapdh was used as a reference protein. Data are averages of three independent experiments.

mTOR levels were significantly upregulated in N2a-APPswe cells by treatments with 20 μM Hcy-thiolactone (1.54±0.20) or 10 μM *N*-Hcy-protein (1.42±0.14), compared to untreated cells (1.00±0.09, *P* < 0.004; **Figure 2C**). Levels of pmTOR were also significantly upregulated by these treatments: 20 μM Hcy-thiolactone (1.36±0.17) or 10 μM *N*-Hcy-protein (1.46±0.11) compared to control (1.00±0.14, *P* < 0.013; **Figure 2D**). Autophagy-related proteins Bcln1, Atg5, and Atg7 were significantly downregulated (by 25-50%, *P* < 0.002) in cells treated with Hcy-thiolactone or *N*-Hcy-protein (**Figure 2E, F,** and **2G**, respectively).

APP levels were significantly upregulated in N2a-APPswe cells treated with 20 μM Hcy-thiolactone (1.74-fold, *P* = 0.001) or *N*-Hcy-protein (1.32-fold, *P* = 0.013) compared to untreated controls (**Figure 2H**).

Similar effects were observed in N2a-APPswe cells treated with 10-fold higher concentrations of Hcy-thiolactone (200 μM), or 2-fold higher *N*-Hcy-protein (20 μM) (**Figure 2**). These findings show that Hcy-thiolactone and *N*-Hcy-protein, which are elevated in *Blmh*^-/-^ mice (4), can affect the expression of Phf8 and its downstream targets.

### 3.6 Blmh depletion by RNA interference increases H4K20me1 biding to mTOR promoter in N2a-APPswe cells

To determine whether increased levels of the histone H4K20me1 mark can promote mTOR gene expression by binding to its promoter in Blmh-depleted cells, we carried out chromatin immunoprecipitation (ChIP) experiments using anti-H4K20me1 antibody (**Figure 3**). The *Blmh* gene was silenced by transfecting N2a-APPswe cells using two different *Blmh*-targeting siRNAs. The cells were permeabilized, treated with anti-H4K20me1 antibody and a recombinant micrococcal nuclease-protein A/G. DNA fragments released form N2a-APPswe cells were quantified by RT-qPCR using primers targeting the transcription start site (TSS) of the mTOR gene as well as upstream (UP) and downstream (DOWN) regions from the TSS.

**Figure 3.**
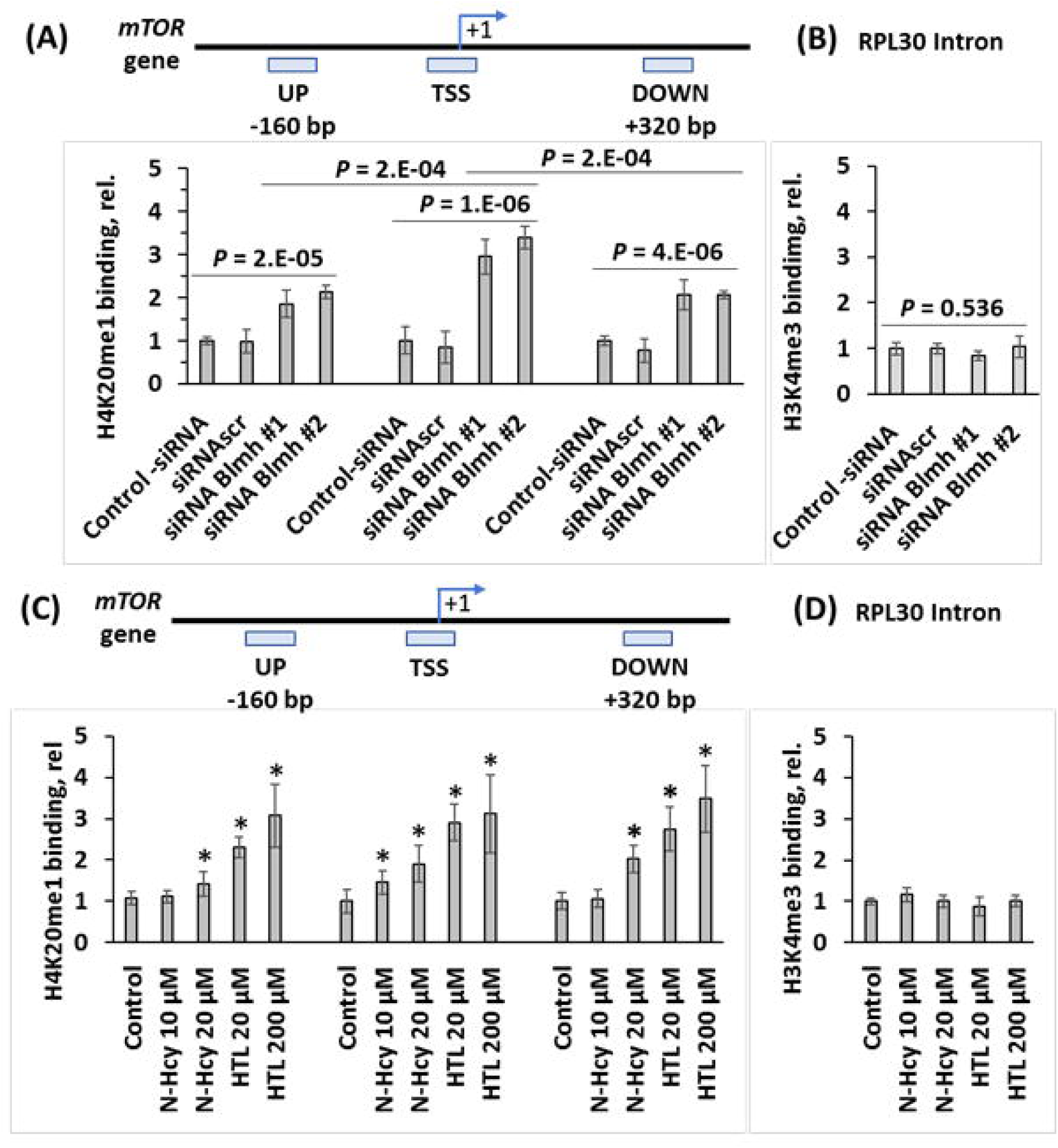
Blmh depletion or treatment with Hcy-thiolactone or *N*-Hcy-protein increases H4K20me1 binding at the *mTOR* promoter in mouse neuroblastoma N2a-APPswe cells. (**A**) CHIP assays with anti-H4K20me1 antibody show the specific binding of H4K20me1 at the transcription start site (TSS) of the *mTOR* gene as well as downstream and upstream sites in Blmh siRNA-silenced N2a-APPswe cells. Bar graphs show the relative H4K20me1 binding at indicated regions of the *mTOR* gene in N2a-APPswe cells transfected with two different siRNAs targeting the *Blmh* gene (siRNA *Blmh* #1 and #2). Transfections without siRNA (Control -siRNA) or with scrambled siRNA (siRNAscr) were used as controls. (**B**) Control CHIP experiment with anti-H3K4me3 antibody shows that *Blmh* gene-silencing did not affect the binding of H3K4me3 at the Rpl30 intron. RT-qPCR was carried out on the input and precipitated DNA fragments. (**C**) N2a-APPswe cells were treated with indicated concentrations of *N*-Hcy-protein, Hcy-thiolactone (HTL), or Hcy for 24h at 37^◦^C. Untreated cells were used as controls. CHIP assays with anti-H4K20me1 antibody show the binding of H4K20me1 at the transcription start site (TSS) of the *mTOR* gene as well as downstream and upstream sites. Bar graphs show the relative H4K20me1 binding at the indicated regions of the *mTOR* gene. (**D**) Control CHIP experiment with anti-H3K4me3 antibody shows that Hcy-thiolactone, *N*-Hcy-protein, and Hcy did not affect the binding of H3K4me3 at the Rpl30 intron. Panels (**C**) and (**D**) were reproduced with permission from ref. (45). RT-qPCR was carried out on the input and precipitated DNA fragments. Data are averages of three independent experiments. * Significant difference *vs*. control, *P* < 0.05.

We found that, in *Blmh*-silenced N2a-APPswe cells, the binding of H4K20me1 was significantly increased at the mTOR TSS (2.9 to 3.4-fold, *P* = 1.E-06), mTOR UP (1.9 to 2.1-fold, *P* = 2.E-05), and mTOR DOWN (2.0-fold, *P* = 4.E-06) sites (**Figure 3A**).

Importantly, in *Blmh*-silenced cells we found significantly more DNA fragments from the mTOR TSS site (2.9±0.4 and 3.4±0.3 for siRNA *Blmh* #1 and #2, respectively) than from the mTOR UP (1.9±0.3 and 2.1±0.2 for siRNA *Blmh* #1 and #2, respectively; *P* = 2.E-04) and mTOR DOWN sites (2.1±0.4 and 2.1±0.1 for siRNA *Blmh* #1 and #2, respectively; *P* = 2.E-04) (**Figure 3A**). Control experiments showed that binding of H3K4me3 to RPL30 intron was not affected by *Blmh* gene silencing (**Figure 3B**). These findings indicate that Blmh depletion induced H4K20me1 binding at the *mTOR* gene, significantly higher at mTOR TSS than at UP and DOWN sites in the Blmh-depleted cells.

CHIP experiments using anti-Phf8 antibody showed that Blmh depletion did not affect binding of Phf8 at the mTOR gene (not shown).

### 3.7 Hcy-thiolactone and *N*-Hcy-protein increase H4K20me1 binding to mTOR promoter in N2a-APPswe cells

Because treatments with Hcy-thiolactone or *N*-Hcy-protein, metabolites that are elevated in Blmh-depleted mice (4), upregulated mTOR expression (**Figure 2C**), it is likely that each of these metabolites can influence mTOR expression by promoting H4K20me1 binding at its promoter. To test this contention, we carried out CHIP experiments with N2a-APPswe cells treated with Hcy-thiolactone or *N*-Hcy-protein using anti-H4K20me1 antibody and determined the extent of H4K20me1 binding at the *mTOR* gene.

We found that Hcy-thiolactone at 20 μM significantly increased binding of H4K20me1 at the *mTOR* TSS (2.9-fold, *P* = 0.003), UP (2.3-fold, *P* = 0.015), and DOWN (2.8-fold, *P* = 0.038) sites. Similar results were obtained with 200 μM Hcy-thiolactone (45) (**Figure 3C**).

*N*-Hcy-protein at 20 μM significantly increased binding of H4K20me1 at the *mTOR* TSS (1.9-fold, *P* = 0.040) and DOWN (2.0-fold, *P* = 0.010) sites, but not UP site (1.4-fold, *P* = 0.089). Smaller, nonsignificant increases were observed with 10 μM *N*-Hcy-protein (45). Control experiments show that the binding of H3K4me3 to RPL30 intron was not affected by treatments with Hcy-thiolactone or *N*-Hcy-protein (45) (**Figure 3D**). Similar results were obtained with N2a cells (not shown).

Binding of Phf8 at the *mTOR* TSS, UP and down sites was not affected by Hcy-thiolactone or *N*-Hcy-protein (not shown).

### 3.8 Blmh depletion by RNA interference or treatments with Hcy-thiolactone and *N*-Hcy-protein promotes Aβ accumulation in N2a-APPswe cells

To determine whether Blmh depletion affects Aβ accumulation, we silenced the *Blmh* gene in N2a-APPswe cells by using RNA interference and quantified Aβ by fluorescence confocal microscopy using anti-Aβ antibody. Towards this end, we transfected N2a-APPswe cells using two different *Blmh*-targeting siRNAs, the cells were permeabilized, treated with anti-Aβ antibody, Aβ was visualized with fluorescent secondary antibody (**Figure 4A**) and quantified (**Figure 4B**). We found that Blmh-silencing (**Figure S2A**) led to increased Aβ accumulation (**Figure 4B**), manifested by significant increases in area (from 133±18 and 130±7 μm^2^ for control -siRNA and siRNA scramble, respectively, to 188±14 μm^2^ and 192±16 μm^2^ for siRNA Blmh #1 and #2, respectively; *P* = 0.007) and average size (from 0.38±0.02 and 0.32±0.02 for control -siRNA and siRNA scramble, respectively, to 0.86±0.06 μm^2^ and 0.86±0.01 μm^2^ for siRNA Blmh #1 and #2, respectively; *P* = 1.E-07) of fluorescent Aβ puncta in Blmh siRNA-treated N2a-APPswe cells compared to siRNAscr-treated or control without siRNA cells (**Figure 4B**).

**Figure 4.**
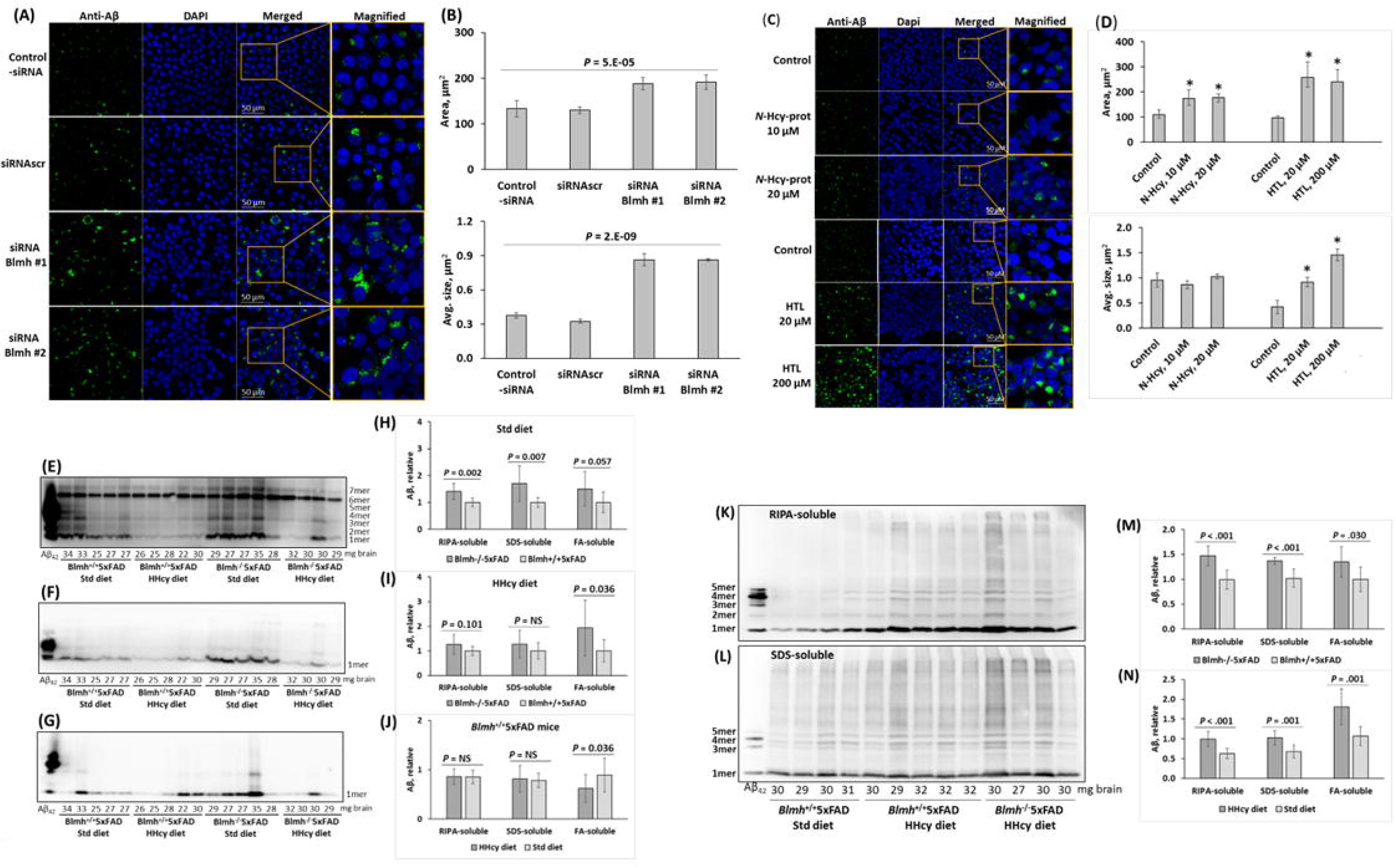
Blmh depletion or treatment with Hcy-thiolactone or *N*-Hcy-protein promotes Aβ accumulation in neural cells and mouse brain. (**A-D**) Analysis of Aβ in mouse neuroblastoma N2a-APPswe cells by confocal immunofluorescence microscopy using anti-Aβ antibody. (**A**, **B**) The cells were transfected with siRNAs targeting the *Blmh* gene (siRNA Blmh #1 and #2). Transfections without siRNA (Control -siRNA) or with scrambled siRNA (siRNAscr) were used as controls. Confocal microscopy images (**A**) and quantification of Aβ signals (**B**) from *Blmh*-silenced and control cells are shown. (**C**, **D**) N2a-APPswe cells were treated with indicated concentrations of *N*-Hcy-protein or Hcy-thiolactone (HTL) for 24h at 37^◦^C. Untreated cells were used as controls. Confocal microscopy images (**C**) and quantification of Aβ signals (**D**) from cells treated with *N*-Hcy-protein or HTL and untreated cells are shown. Each data point is an average of three independent experiments with triplicate measurements in each. Panels (**C**) and (**D**) were reproduced with permission from ref. (45). (**E-N**) Analysis of Aβ in brains from 5-month-old (**E-J**) and 12-month-old (**K-N**) *Blmh*^-/-^5xFAD and *Blmh^+^*^/+^5xFAD mice fed with a HHcy or standard diet since weaning at the age of one month. Brain extracts were analyzed on SDS-PAGE gels and Aβ was quantified by Western blotting. Representative pictures of Western blots of Aβ fractions extracted from brains with RIPA buffer (**E, K**), 2% SDS (**F**, **L**), and 70% formic acid (FA) (**G**) are shown. Numbers below each lane refer to the amount of brain (mg) from each mouse used in the experiment. A commercial Aβ_42_ standard is shown in the first lane from left in each blot. Total Aβ signals in each lane (representing an individual mouse) were determined by scanning of all chemiluminescent bands. Bar graphs (**H**, **I**, **J**, **M**, **N**) show brain Aβ quantification for *Blmh*^-/-^5xFAD and *Blmh^+^*^/+^5xFAD mice (n = 8-10 mice/group). Data points for each mouse represent averages of four independent measurements.

Because Blmh depletion elevates Hcy-thiolactone and *N*-Hcy-protein in mice (4) we examined whether each of these metabolites can induce Aβ accumulation in N2a-APPswe cells. We found significantly more Aβ in cells treated with Hcy-thiolactone (20 - 200 μM) or *N*-Hcy-protein (10 - 20 μM), manifested by significantly increased area of fluorescent Aβ puncta observed in confocal immunofluorescence images, compared to control-siRNA and siRNAscr (**Figure 4C, D**). However, while treatments with Hcy-thiolactone led to increased size and signal intensity of the fluorescent Aβ puncta, treatments with *N*-Hcy-protein did not (**Figure 4D**), suggesting different effects of these metabolites on the structure of Aβ aggregates. These findings suggest that Hcy-thiolactone and *N*-Hcy-protein can contribute to increased accumulation of Aβ induced by Blmh depletion.

### 3.9 Blmh depletion increases Aβ accumulation in brains of 5xFAD mice

To examine if Blmh depletion can promote Aβ accumulation in mouse brain, we generated *Blmh*^-/-^5xFAD mice and their *Blmh^+^*^/+^5xFAD siblings by crossing *Blmh*^-/-^ mice with Aβ-overproducing 5xFAD animals. We prepared SDS-soluble and formic acid (FA)-soluble Aβ fractions, which contain most of the total Aβ (38), as well as a minor Aβ fraction extractable with a RIPA buffer from the brains of 5- and 12-month-old mice fed with a standard chow diet or with HHcy diet (1% Met in drinking water) since weaning at the age of 1 month. Aβ was quantified in these brain extracts by Western blotting using monoclonal anti-Aβ antibody. Representative Western blots are shown in **Figure 4E, F, G, K, L**.

We found that RIPA- and SDS-soluble Aβ were significantly elevated (*P* = 0.002 and 0.007, respectively), and FA-soluble Aβ tended to be elevated (*P* = 0.057) in brains of 5-month-old *Blmh*^-/-^5xFAD mice compared to *Blmh^+^*^/+^5xFAD sibling controls in mice fed with a standard diet (**Figure 4H**). HHcy diet attenuated effects of Blmh depletion on RIPA-soluble and SDS-soluble Aβ, while FA-soluble Aβ was significantly upregulated in 5-month- old *Blmh*^-/-^5xFAD mice (**Figure 4I**). HHcy diet did not affect RIPA-, SDS-, and FA-soluble Aβ in 5-month-old *Blmh*^-/-^5xFAD mice (quantification not shown). In *Blmh^+^*^/+^5xFAD mice, HHcy diet did not affect RIPA-soluble and SDS-soluble Aβ levels, but increased FA-soluble Aβ (**Figure 4J**).

In in brains of 12-month-old HHcy *Blmh*^-/-^5xFAD mice, RIPA-, SDS-, and FA-soluble Aβ were significantly elevated compared with HHcy *Blmh^+^*^/+^5xFAD sibling controls (**Figure 4M**). HHcy diet significantly elevated RIPA-, SDS-, and FA-soluble Aβ in 12-month-old *Blmh*^-/-^5xFAD mice (quantification not shown) RIPA-soluble Aβ was also elevated in brains of HHcy *Blmh^+^*^/+^5xFAD mice (**Figure 4M**).

### 3.10 Phf8 depletion upregulates Aβ but not App in N2a-APPswe cells

The findings that Phf8 expression was significantly reduced in brains of *Blmh*^-/-^ mice (**Figure 1A**) and in Blmh-silenced (**Figure 2B**) or Hcy metabolite-treated (**Figure 2A**) mouse neuroblastoma cells, suggested that Phf8 depletion by itself can affect biochemical pathways leading to Aβ accumulation. To examine this, we depleted Phf8 in N2a-APPswe cells by RNA interference using Phf8-targeting siRNAs and quantified by Western blotting proteins that we found to be affected in the *Blmh*^-/-^ mouse brain (**Figure 1**). We found significantly reduced Phf8 levels (by 80%, *P* = 7.E-09; **Figure S4A**), significantly increased H4K20me1 (3-fold, *P* = 7.E-09; **Figure S4B**), mTOR (1.4-fold, *P* = 7.E-06; **Figure S4C**), and pmTOR (1.6-fold, *P* = 5.E-04; **Figure S4D**) levels in Phf8-silenced cells. Autophagy-related proteins Atg5 and Atg7 were significantly downregulated (by 20-35%, *P* = 3.E-04 – 2.E-06; **Figure S4E, F**) while Bcln1 was not affected in Phf8-silenced cells (**Figure S4G**).

Notably, APP levels were not affected in Phf8-silenced cells (**Figure S4H**). We also quantified Aβ by fluorescence confocal microscopy and found that Aβ was upregulated in Phf8-silenced cells, manifested by significantly increased average size and signal intensity of fluorescent Aβ puncta compared to controls without siRNA or with siRNAscr (**Figure S4I, J**). These findings suggest that upregulation of Aβ in Phf8-silenced cells was caused by impaired autophagy and not by increase in APP levels.

### 3.11 Blmh depletion impairs recognition memory, induces neurodegeneration, and sensorimotor deficits

To examine effects of Blmh depletion on cognition and sensorimotor activity, *Blmh*^-/-^ and *Blmh*^+/+^ mice fed with a control or HHcy diet were assessed in the novel object recognition (NOR), hindlimb clasping, and ledge tests. We found that 4-month-old *Blmh*^-/-^ mice did not differentiate between novel and familiar objects in the NOR test, regardless of the diet (**Figure 5A, B**), indicating impaired recognition memory. As expected, 4-month-old *Blmh*^+/+^ mice showed normal preference for novelty (*P* = 0.001); however, the preference for novelty disappeared when the mice were fed with a HHcy diet (*P* = NS). In contrast, 2-month-old *Blmh*^-/-^ mice showed significant preference for a novel object, regardless of the diet (*P* = 2.E-09 to 1.E-04), as did 2-month-old *Blmh*^+/+^ mice (*P* = 0.002). These findings indicate that a longer exposure (>2-month) is required for the manifestation of detrimental effects of Blmh depletion or HHcy on cognition.

**Figure 5.**
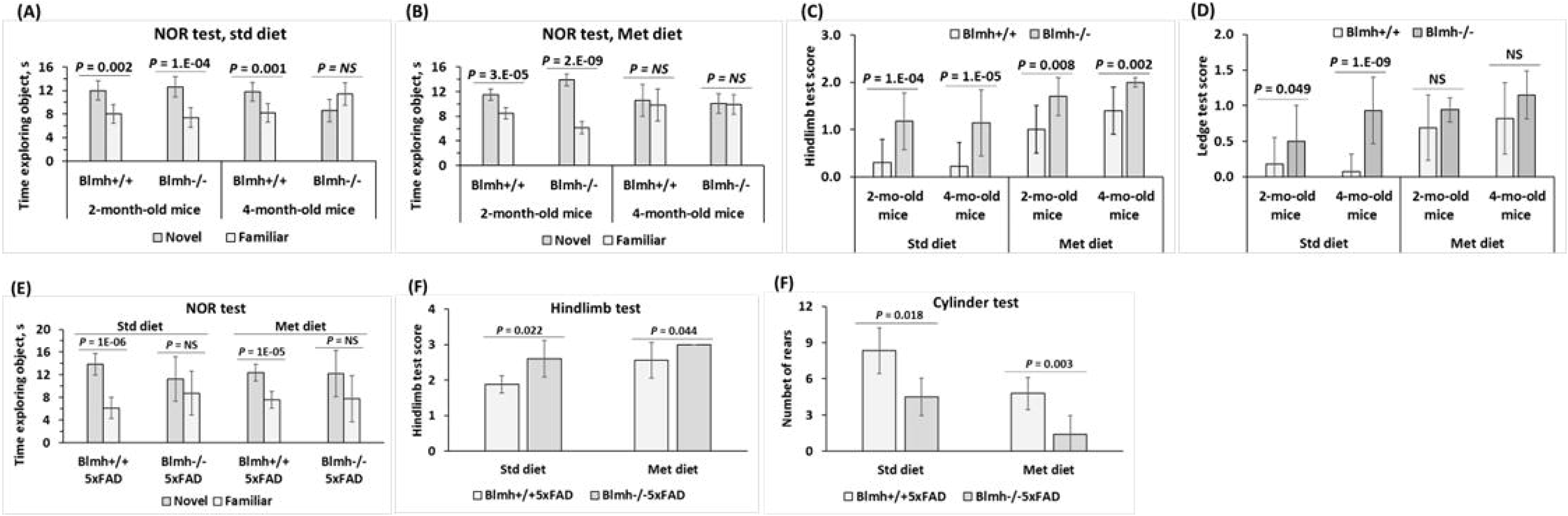
Blmh depletion impairs recognition memory and sensorimotor activity. *Blmh*^-/-^ *vs*. *Blmh*^+/+^ mice: (**A, B**) Novel object recognition test: Time spent at novel and familiar objects. N = 6, 7, 9, 4 mice/group and 7, 7, 8, 10 mice/group. (**C**) Hindlimb clasping test scores. N = 8, 9, 30, 14 mice/group. (**D**) Ledge test scores. N = 8, 9, 30, 14 mice/group. *Blmh*^-/-^5xFAD *vs*. *Blmh*^+/+^5xFAD mice: (**E**) Novel object recognition test: Time spent at novel and familiar objects. N = 8, 4, 8, 5 mice/group. (**F**) Hindlimb clasping test scores. N = 8, 4, 9, 7 mice/group. (**G**) Cylinder test: number of rears. N = 8, 6, 9, 5 mice/group.

The hindlimb test showed more severe clasping (significantly higher scores) in *Blmh*^-/-^ mice *vs*. their *Blmh*^+/+^ littermates (2-month-old: 1.2±0.6 *vs*. 0.3±0.5, *P*_genotype_ = 1.E-04; 4-month-old: 1.1±0.7 *vs*. 0.2±0.5, *P*_genotype_ = 1.E-05; **Figure 5C**), indicating neurodegeneration in *Blmh*^-/-^ animals. HHcy diet significantly increased the hindlimb scores, with greater increases in *Blmh*^+/+^ (2-month-old: from 0.3±0.5 to 1.0±0.5, *P*_diet_ = 0.002; 4-mo-old: from 0.2±0.5 to 1.4±0.5, *P*_diet_ = 1.E-08) than in *Blmh*^-/-^ mice (2-mo-old: from 1.2±0.6 to 1.7±0.4, *P*_diet_ = 0.038; 4-mo-old: from 1.1±0.7 to 2.0±0.0, *P*_diet_ = 0.001), resulting in attenuated *Blmh*^-/-^ vs. *Blmh*^+/+^ difference in HHcy animals (**Figure 5C**). Overall, HHcy diet attenuated effects of the *Blmh*^-/-^ on the hindlimb score.

The ledge test showed significantly higher scores for *Blmh*^-/-^ vs. *Blmh*^+/+^ mice (2-month-old: 0.5±0.5 *vs*. 0.2±0.4, *P*_genotype_ = 0.049; 4-month-old: 0.93±0.47 *vs*. 0.07±0.25, *P*_genotype_ = 1.E-09; Figure XD), indicating neuromotor deficiency in *Blmh*^-/-^ animals. HHcy diet increased the ledge test scores, with greater increases in *Blmh*^+/+^ (2-month-old: from 0.2±0.4 to 0.7±0.5, *P*_diet_ = 0.005; 4-mo-old: from 0.07±0.25 to 0.8±0.5, *P*_diet_ = 5.E-08) than in *Blmh*^-/-^ mice (2-month-old: from 0.5±0.5 to 0.9±0.2, *P*_diet_ = 0.020; 4-mo-old: from 0.93±0.47 to 1.15±0.34, *P*_diet_ = 0.087), resulting in attenuated *Blmh*^-/-^ vs. *Blmh*^+/+^ difference in HHcy animals (**Figure 5D**). Overall, HHcy diet attenuated effects of the *Blmh*^-/-^ on the ledge test score.

We also examined effects of Blmh depletion on cognition and neuromotor activity in *Blmh*^-/-^5xFAD *vs*. *Blmh*^+/+^5xFAD mice fed with a standard or HHcy chow diet. We found that 1-year-old *Blmh*^-/-^5xFAD mice did not differentiate between novel and familiar objects in the NOR test, regardless of the diet (*P* = NS, **Figure 5E**), indicating impaired recognition memory in these animals. In contrast, 1-year-old *Blmh*^+/+^5xFAD mice fed with a standard chow diet or HHcy diet showed normal preference for novelty (*P* = 1.E-06 or 1E.-05, respectively).

The hindlimb clasping test showed significantly higher scores in *Blmh*^-/-^5xFAD *vs*. *Blmh*^+/+^5xFAD mice fed with a standard chow or HHcy diet (standard diet: 3.0±0.0 *vs*. 2.6±0.5, *P*_genotype_ = 0.044; HHcy diet: 2.6±0.5 *vs*. 1.9±0.2, *P*_genotype_ = 0.022; **Figure 5F**), indicating that *Blmh*^-/-^ genotype promotes neurodegeneration in *Blmh*^-/-^5xFAD animals. HHcy diet increased the hindlimb scores, with greater increases in *Blmh*^+/+^5xFAD (from 1.9±0.2 to 2.6±0.5, *P*_diet_ = 0.034) than in *Blmh*^-/-^ mice (from 2.6±0.5 to 3.0±0.0, *P*_diet_ = 0.079), resulting in attenuated *Blmh*^-/-^5xFAD vs. *Blmh*^+/+^5xFAD difference in HHcy animals (**Figure 5F**).

The ledge test showed similar scores for *Blmh*^-/-^5xFAD *vs*. *Blmh*^+/+^5xFAD mice fed with a standard chow or HHcy diet (standard diet: 2.6±0.5 *vs*. 2.2±0.4, *P*_genotype_ = 0.329; HHcy diet: 3.0±0.0 *vs*. 2.7±0.4, *P*_genotype_ = 0.120). HHcy diet increased the ledge test scores in *Blmh*^+/+^5xFAD (from 2.2±0.4 to 2.7±0.4, *P*_diet_ = 0.114) and *Blmh*^-/-^5xFAD mice (from 2.6±0.5 to 3.0±0.0, *P*_diet_ = 0.037).

The cylinder test showed significantly reduced scores for *Blmh*^-/-^5xFAD *vs*. *Blmh*^+/+^5xFAD mice fed with a standard chow or HHcy diet (standard diet: 4.5±1.9 *vs*. 8.3±3.3, *P*_genotype_ = 0.018; HHcy diet: 1.4±1.3 *vs*. 4.8±1.7, *P*_genotype_ = 0.003; **Figure 5G**), indicating that *Blmh*^-/-^ genotype promotes neurodegeneration in *Blmh*^-/-^5xFAD animals. HHcy diet reduced the beaker test scores, with greater reductions in *Blmh*^-/-^5xFAD (from 4.5±1.9 to 1.4±1.3, *P*_diet_ = 0.010) than *Blmh*^+/+^5xFAD mice (from 8.3±3.3 to 4.5±1.9, *P*_diet_ = 0.001), resulting in increased *Blmh*^-/-^5xFAD *vs*. *Blmh*^+/+^5xFAD difference in HHcy animals (**Figure 5G**).

## 4 DISCUSSION

Previous findings showed that Blmh activity/expression was reduced in brains from Alzheimer’s disease patients (29). In mice, Blmh depletion elevated brain Hcy-thiolactone levels (4), increased the animals’ susceptibility to Hcy-thiolactone-induced seizures (4), caused astrogliosis (30), and pro-neurodegenerative changes in brain proteome (8). These findings suggest that Blmh plays an important protective role in the CNS.

Our present findings show that Blmh1 protects from amyloidogenic APP processing to Aβ and unravel the mechanistic basis of the protective role of Blmh in the CNS. Specifically, we found that genetic Blmh depletion in the mouse brain and mouse neuroblastoma cells downregulated histone demethylase Phf8 and upregulated H4K20me1 epigenetic mark. This led to mTOR signaling upregulation, autophagy downregulation, and resulted in Aβ upregulation. Measurements of protein and mRNA levels indicated that Blmh depletion dysregulated the Phf8-> mTOR->autophagy pathway at the transcriptional level.

In previous studies we found that Blmh is a Hcy-thiolactone-hydrolyzing enzyme (3) and that Hcy-thiolactone and *N*-Hcy-protein were elevated in *Blmh*^-/-^ mice (4). In the present study we found that Blmh depletion or treatments with Hcy-thiolactone and *N*-Hcy-protein increased H4K20me1 binding at the mTOR promotor (**Figure 3**) and upregulated mTOR signaling (**Figure 1C, D**; **Figure 2C, D**). This indicates that Blmh is a negative regulator of mTOR signaling by controlling levels of Hcy metabolites that affect occupancy of the mTOR promotor by H4K20me1. The effects of Hcy-thiolactone and *N*-Hcy-protein on mTOR signaling can be explained by our findings showing that Phf8, the epigenetic regulator of mTOR expression, was downregulated by these metabolites (**Figure 2A**) *and* by Blmh depletion (**Figure 1A, Figure S2B, Figure S3B**) whereas the histone H4K20me1 mark was upregulated (**Figure 1B, 2B; Figure S2C**). These findings provide direct mechanistic evidence linking Hcy-thiolactone and *N*-Hcy-protein with dysregulated mTOR signaling and its downstream consequences such as upregulation of Aβ due to impaired autophagy. This mechanism is further supported by our findings showing that effects of Phf8 silencing by RNA interference on mTOR, autophagy, and Aβ (**Figure S4**) were akin to the effects of treatments with Hcy-thiolactone or *N*-Hcy-protein (**Figure 2A**).

In the present study we found that depletion of Blmh upregulated APP (**Figure 1H; Figure S1H, Figure S2I, Figure S3G**). In contrast, depletion of Phf8 did not affect APP expression (**Figure S4G**). These findings suggest that Blmh interacts with APP in the mouse brain while Phf8 does not. The Blmh-APP interaction is most likely direct, as suggested by findings of other investigators. For example, one study has shown that human BLMH interacts with APP *in vitro* and that overexpressed BLMH has the ability to process human APP to Aβ in the 293-HEK and CHO cells (20). Another study reported that rat Blmh can further hydrolyze Aβ *in vitro*, with fibrillar Aβ40 and Aβ42 being more resistant than nonfibrillar peptides (46).

Although Blmh depletion downregulated Phf8 and upregulated APP (**Figure 1H, Figure S1H, Figure S2I, Figure S3G**) and Aβ (**Figure 4A, B, E, F**), depletion of Phf8 upregulated Aβ (**Figure S4I, J**) but had no effect of APP (**Figure S4G**) in mouse neuroblastoma cells. These findings suggest that two pathways can lead to increased Aβ generation in Blmh-depleted brain and neural cells. One pathway involves Hcy metabolites, which upregulate APP while another pathway involves impaired Aβ clearance due to downregulated autophagy. Only one mechanism, downregulated autophagy, lead to accumulation of Aβ in Phf8-depleted cells (**Figure 6**).

**Figure 6.**
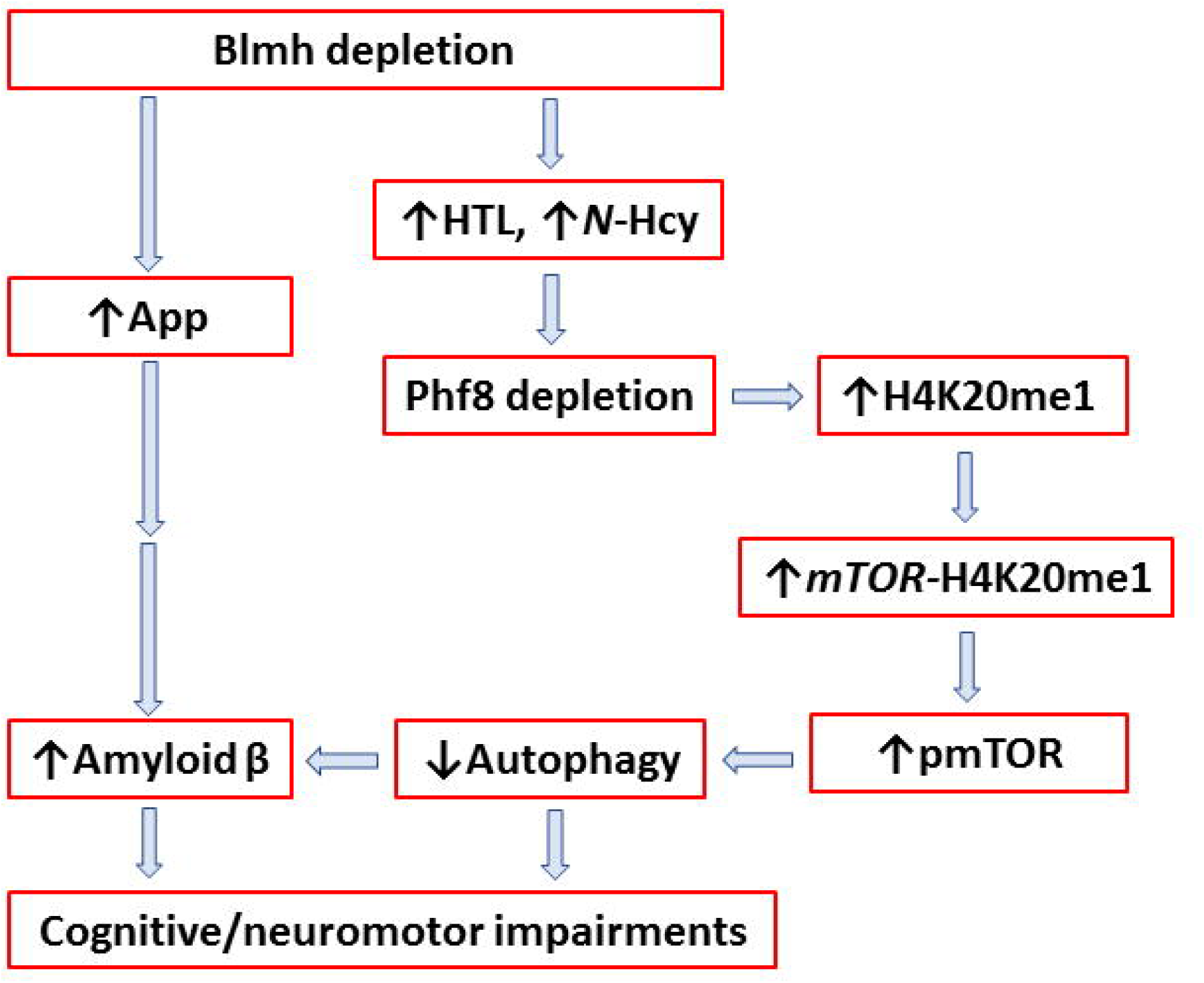
Mechanisms underlying Aβ generation in Blmh-depleted brains. Up and down arrows show direction of changes in the indicated metabolites, proteins, and molecular processes. Blmh, bleomycin hydrolase; Hcy, homocysteine; HTL, Hcy-thiolactone; APP, amyloid beta precursor protein; mTOR, mammalian target of rapamycin; pmTOR, phospho-mTOR; Phf8, Plant Homeodomain Finger protein 8.

5xFAD mice accumulate high levels of Aβ beginning around 2 months of age (37) and develop cognitive impairments beginning at 4-5 months of age and sensorimotor impairments at about 9 months of age (47) (https://www.alzforum.org/research-models/5xfad-b6sjl). For example, 5xFAD mice perform worse than the wild type animals in the memory and sensorimotor tests. In the present work we found that Blmh depletion aggravated those impairments. Specifically, *Blmh*^-/-^5xFAD mice showed worse performance than *Blmh*^+/+^5xFAD animals in the NOR test, indicating impaired memory, and in the hindlimb and cylinder tests, indicating sensorimotor impairments (**Figure 5E, F, G**).

We found similar memory and sensorimotor impairments also in *Blmh*^-/-^ *vs*. *Blmh*^+/+^ mice, which do not accumulate Aβ (**Figure 5A, B, C, D**). These findings suggest that Blmh depletion can cause memory and sensorimotor impairments in Aβ-dependent and -independent manner.

Memory and sensorimotor impairments in *Blmh*^-/-^ and *Blmh*^-/-^5xFAD mice can be caused, at least in part, by the depletion of Phf8, which we found in the present study. Indeed, in humans PHF8 depletion is linked to intellectual disability, autism spectrum disorder, attention deficit hyperactivity disorder (31), and mental retardation (32), while similar neurological deficits were found in *Phf8*^-/-^ mice memory (33). However, PHF8 was not known to be associated with Aβ, a hallmark of Alzheimer’s disease. Our present findings that Phf8 depletion in mouse neuroblastoma cells, induced by *Phf8* siRNA interference (**Figure 5A**), or by supplementation with Hcy-thiolactone, *N*-Hcy-protein, or Hcy (**Figure 2A**), significantly increased Aβ accumulation (**Figure 5I, J**), suggest that Phf8 depletion can also underly the association of Alzheimer’s disease with HHcy.

We also found that HHcy, induced in mice by supplementation of their drinking water with 1% Met, did not change effects of Blmh depletion on mTOR signaling and autophagy in young *Blmh*^-/-^ (**Figure 1**) and *Blmh*^-/-^ 5xFAD mice (2- and 5-month-old) (**Figure S1**). However, in old *Blmh*^-/-^5xFAD mice (12-month) HHcy eliminated effects of Blmh depletion on mTOR signaling and one of the autophagy-related proteins (Atg7) but did not affect others (Atg5 and Bcln1) (**Figure S1**). These findings indicate that in young *Blmh*^-/-^5xFAD mice effects of *Blmh*^-/-^ genotype are dominant over the effects of HHcy diet and that old *Blmh*^-/-^5xFAD mice are more sensitive to detrimental effects of HHcy than young mice. These findings also suggest that in old animals the effects of HHcy on autophagy may be, at least in part, independent of its effects on mTOR signaling.

Importantly, Blmh depletion caused changes in the Phf8->H4K20me1->mTOR->autophagy pathway similar to the changes induced by HHcy **(Figure 1, Figure S1)**. Also, Blmh depletion and HHcy both increased Aβ accumulation in the mouse brain (**Figure 4E-N**). Our previous work has shown that a common primary biochemical outcome of Blmh depletion and HHcy was essentially the same: Blmh depletion led to elevation of Hcy-thiolactone and *N*-Hcy-protein in mice (4), as did HHcy (48). As shown in the present work, Blmh depletion by siRNA interference, as well as treatments with Hcy-thiolactone or *N*-Hcy-protein caused similar biochemical outcomes in mouse neuroblastoma cells, *i. e.*, downregulation of Phf8 expression, which then triggered downstream effects on H4K20me1 binding to mTOR (**Figure 3**), autophagy, APP, and Aβ (**Figure 2**). These findings suggest that Aβ accumulation in Blmh-depleted mouse brain is mediated by the effects of Hcy-thiolactone and *N*-Hcy-protein on the Phf8, mTOR, and autophagy (**Figure 6**).

In conclusion, we found that Blmh depletion upregulated mTOR signaling *via* Phf8/H4K20me1, downregulated autophagy, which led to Aβ accumulation and cognitive and neuromotor deficits in mice. Upregulated APP expression in Blmh-depleted mice can also contribute to Aβ accumulation. By revealing a mechanism by which Blmh prevents Aβ generation and cognitive/neuromotor deficits, our findings significantly expand our understanding how Blmh maintains brain homeostasis.

## AUTHOR CONTRIBUTIONS

Ł. Witucki performed and analyzed the experiments; Ł. Witucki and K. Borowczyk, carried out behavioral assessments; J. Suszyńska-Zajczyk analyzed NOR data; P. Pawlak and E. Warzych provided advice on confocal microscopy experiments; H. Jakubowski conceived the idea for the project, designed the study, generated *Blmh*^-/-^5xFAD mouse model, bred the mice, collected tissue samples, analyzed data, and wrote the paper.

## ACKNOWLEDGMENTS

We thank S. S. Sisodia for kindly providing mouse neuroblastoma N2a-APPswe cells expressing human APP-695 harboring the K670N/M671L Swedish double mutation associated with familial early-onset Alzheimer’s disease. Supported in part by Grants 2018/29/B/NZ4/00771, 2019/33/B/NZ4/01760, and 2021/43/B/NZ4/00339 from the National Science Center, Poland, and Grant 17GRNT32910002 from the American Heart Association.

## DATA AVAILABILITY STATEMENT

The data that support the findings of this study are available in the methods and/or supplementary material of this article.

