## Supplementary for "Depletion of bleomycin hydrolase (Blmh) downregulates histone demethylase Phf8, impairs mTOR signaling/autophagy, accelerates amyloid beta accumulation, and induces neurological deficits in mice"

**Supplementary Material**

**Supplementary Figures S1, S2, S3, S4**

**Supplementary Table S1**

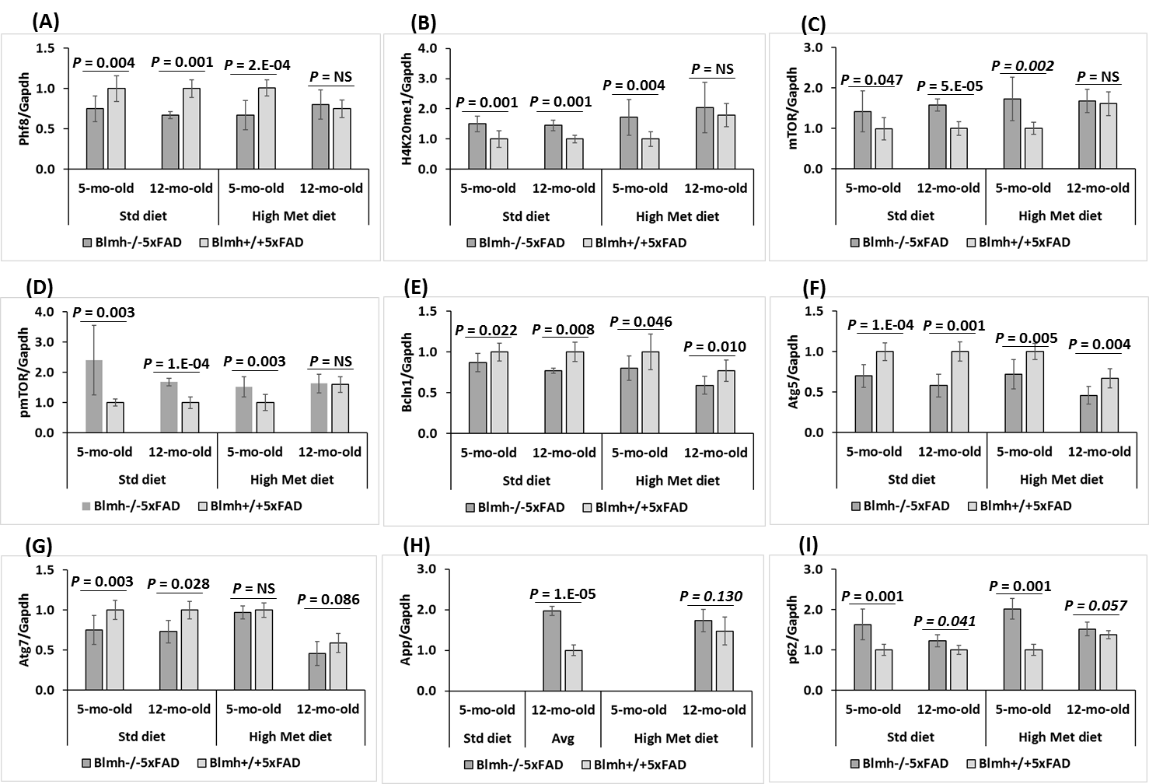

**Figure S1.** Blmh depletion affects the expression of histone demethylase Phf8, histone H4K20me1 epigenetic mark, mTOR signaling, autophagy, and App in the *Blmh*^-/-^5xFAD mouse brain. One-month-old *Blmh*^-/-^5xFAD mice and *Blmh*^+/+^5xFAD sibling controls fed with HHcy high Met diet (1% Met in drinking water) or control diet for 4 and 11 months were used in experiments. Each group included 7 - 10 mice of both sexes. Bar graphs illustrating quantification of the following brain proteins by Western blotting are shown: Phf8 (**A**), H4K20me1 (**B**), mTOR (**C**), pmTOR (**D**), Bcln1 (**E**), Atg5 (**F**), Atg7 (**G**), App (**H**), and p62 (**I**). Gapdh protein were used as references for normalization. Pictures of Western blots used for protein quantification are shown in panel (**J**). Data are averages of three independent experiments.

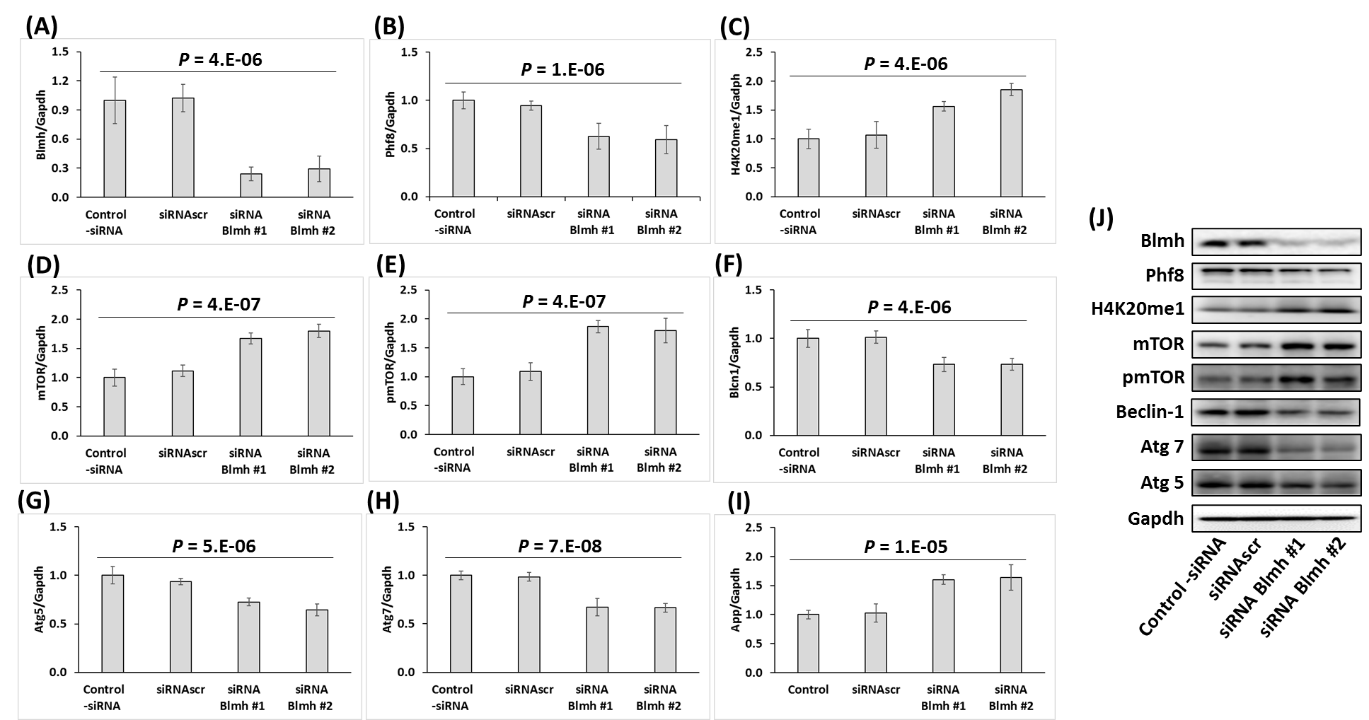

**Figure S2**. *Blmh* gene silencing in mouse neuroblastoma N2a-APPswe cells recapitulates changes in histone demethylase Phf8, H4K20me1, mTOR signaling, APP, and autophagy-related protein levels observed in *Pon1*^-/-^ mouse brain. Bar graphs illustrating the quantification of Blmh (**A**), Phf8 (**B**), H4K20me1 (**C**), mTOR (**D**), pmTOR (**E**), Bcln1 (**F**), Atg5 (**G**), Atg7 (**H**), and App (**I**) in N2a-APPswe cells transfected with two different siRNAs targeting the *Blmh* gene (siRNA *Blmh* #1 and #2) are shown. Transfections without siRNA (Control -siRNA) or with scrambled siRNA (siRNAscr) were used as controls. Representative Western blots are shown in panel (**J**). Gapdh was used as a reference protein. Data are averages of three independent experiments.

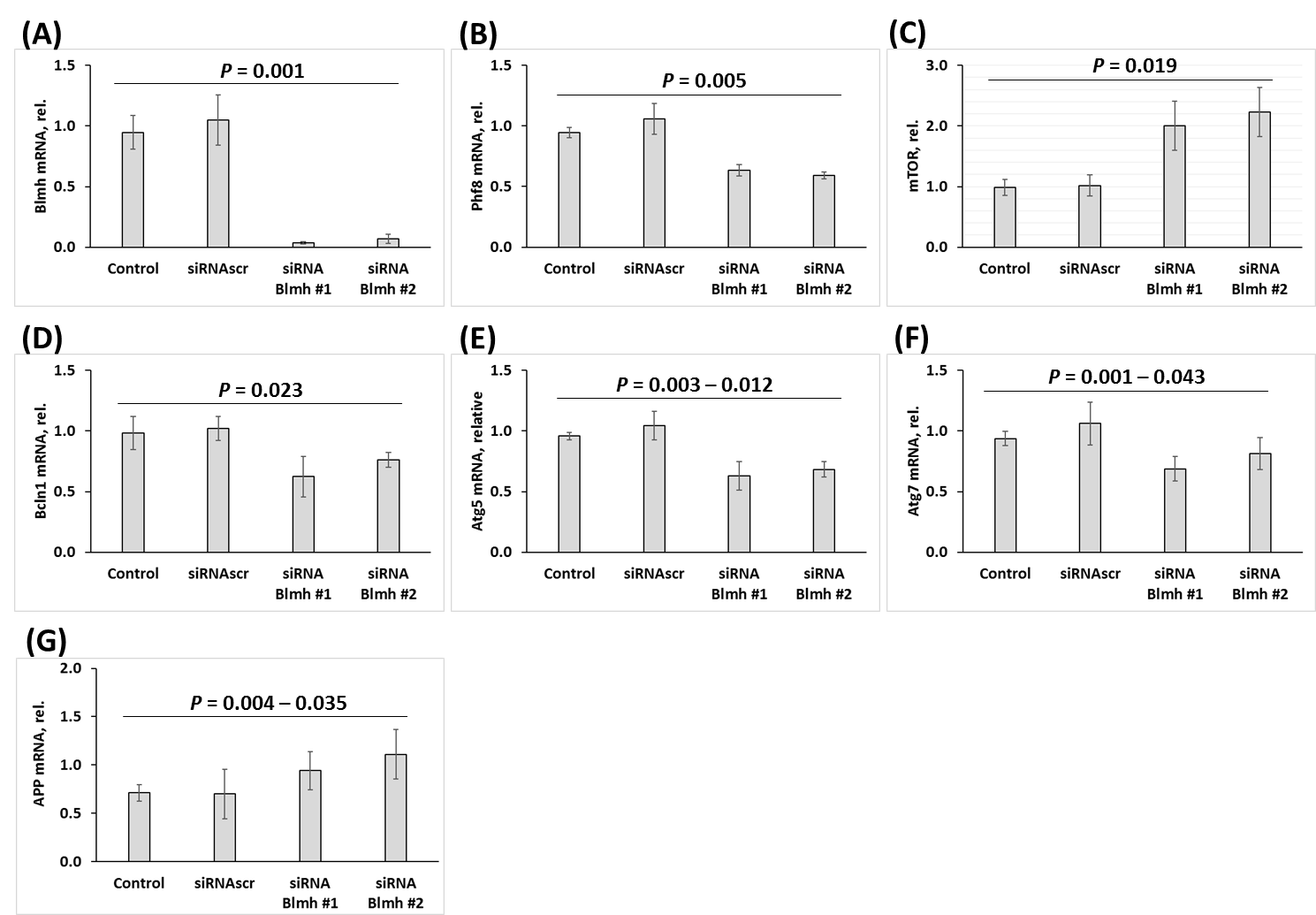

**Figure S3.** *Blmh* gene silencing affects expression of mRNAs for Phf8, mTOR, APP, and autophagy-related proteins in mouse neuroblastoma N2a-APPswe cells. Bar graphs illustrating the quantification by RT-qPCR of mRNAs for Blmh (**A**), Phf8 (**B**), mTOR (**C**), Bcln1 (**D**), Atg5 (**E**), Atg7 (**F**), and App (**G**), in N2a-APPswe cells transfected with two different siRNAs targeting the *Blmh* gene (siRNA Blmh #1 and #2) are shown. Gapdh mRNA was used as a reference. Transfections without siRNA (Control) or with scrambled siRNA (siRNAscr) were used as controls.

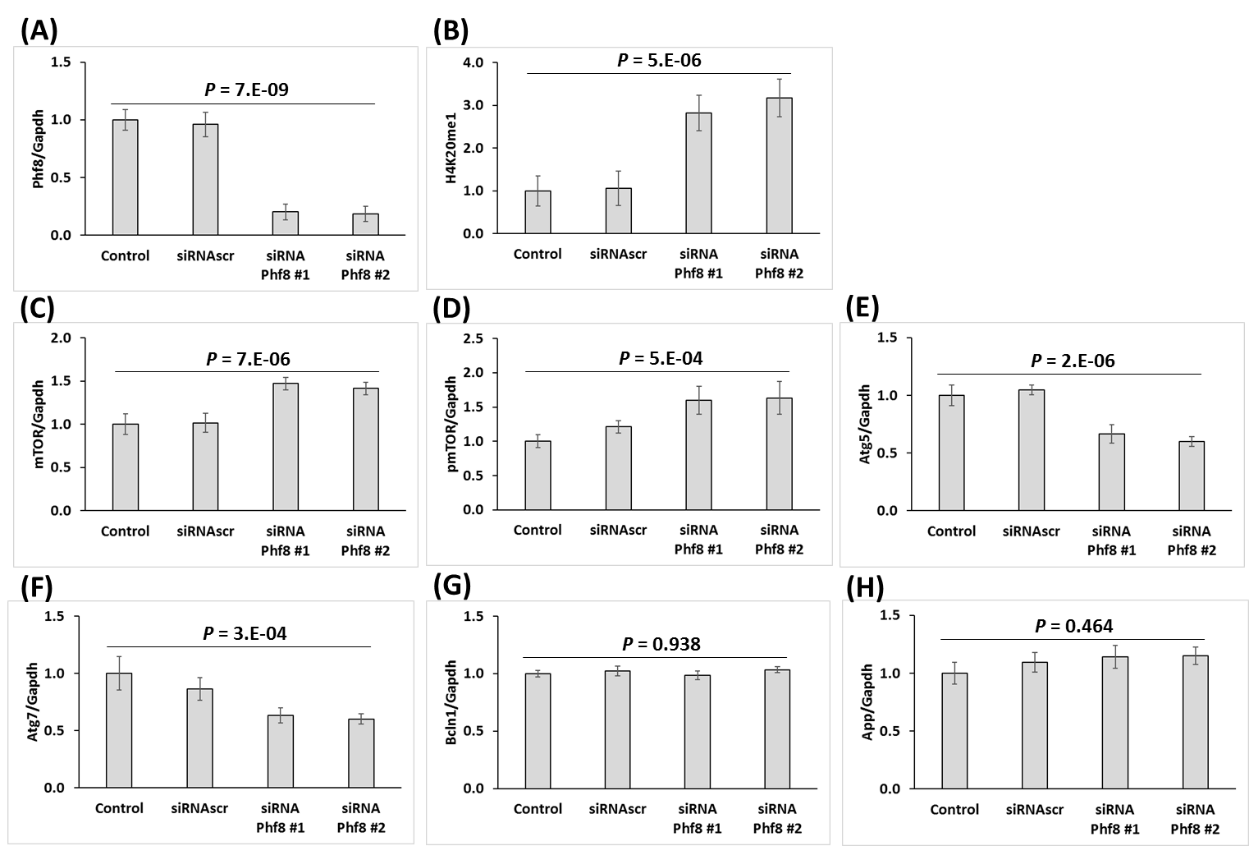

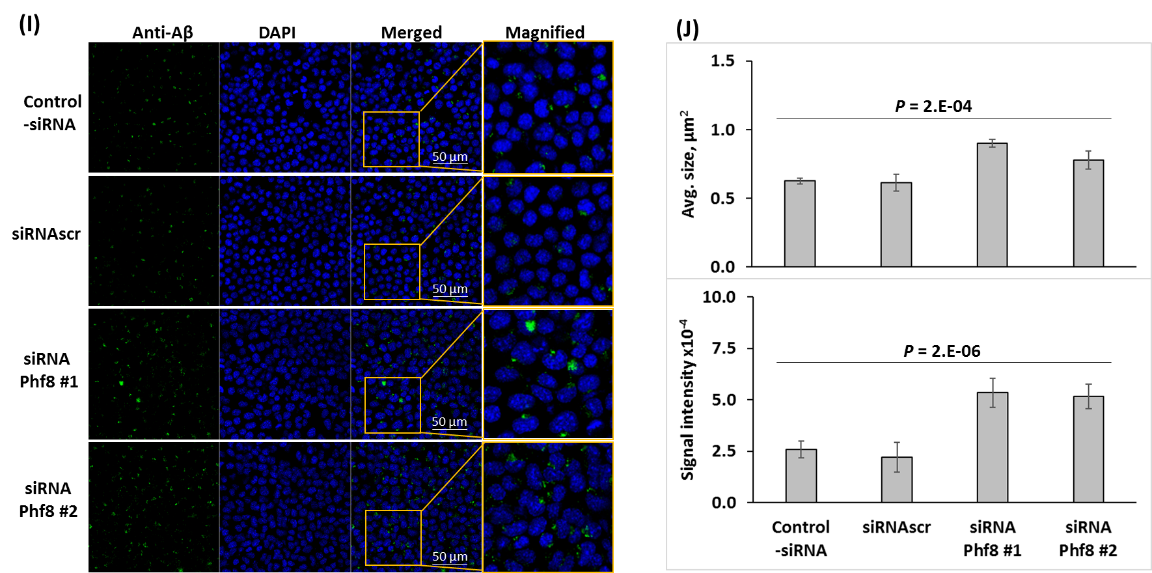

**Figure S4.** Phf8 depletion promotes Aβ accumulation mediated by upregulation of mTOR signaling and inhibition of autophagy in the mouse neuroblastoma N2a-APPswe cells. The cells were transfected with siRNAs targeting the *Phf8* gene (Phf8 siRNA #1 and #2). Transfections without siRNA (Control -siRNA) or with scrambled siRNA (siRNAscr) were used as controls. Proteins were quantified by Western blotting. Bar graphs illustrate levels of **(A)** Phf8, **(B)** H4K20me1, **(C)** mTOR, **(D)** pmTOR, **(E)** Atg5, **(F)** Atg7, **(G)** Bcln1, and **(H)** App. Aβ was detected and quantified by confocal immunofluorescence microscopy using anti-Aβ antibody. (**I**) Confocal microscopy images of Aβ signals from *Phf8*-silenced and control N2a-APPswe cells. (**J**) Bar graphs show quantification of Aβ signals.

| **Table S1 Primers used for PCR or RT-qPCR** | |
| --- | --- |
| **Gene** | **Primer sequence** |
| APP  App | Forward: 5′-CTTCCCCAAGATCCTGATAAACT-3′ |
|  | Reverse: 5′-CCGGGTGTCTCCAGGTACT-3′ |
| Atg5 | Forward: 5′-AAGGCACACCCCTGAAATGG-3′ |
|  | Reverse: 5′-TGATGTTCCAAGGAAGAGCTGAA-3′ |
| Atg7 | Forward: 5′-GCCAACTCCACACTGCTTTC-3′ |
|  | Reverse: 5′-TCTTCTGGGTCAGTTCGTGC-3′ |
| β-actin | Forward: 5′-GCAGGAGTACGATGAGTCCG-3′ |
|  | Reverse: 5′-ACGCAGCTCAGTAACAGTCC-3′ |
| Beclin-1 | Forward: 5′-GAGGAAGCTCAGTACCAG CG-3′ |
| Blmh | Reverse: 5′-CCAGATGTGGAAGGTGGCAT-3′ |
|  | Forward p1: 5′-CACTGTAGCTGTACTCACAC-3’ |
|  | Reverse p2: 5′-GCGACAGAGTACCATGTAGG-3′ (exon 3); Reverse p3: 5′-ATTTGTCACGTCCTGCACGACG-3′ (neomycin cassette) |
| hAPP transgene  in 5xFAD mice | Forward: 5′-AGAGTACCAACTTGCATGACTACG-3′; |
|  | Reverse: 5′-ATGCTGGATAACTGCCTTCTTATC-3′ |
| hPS1 transgene  in 5xFAD mice | Forward: 5′-GCTTTTTCCAGCTCTCATTTACTC-3′ |
|  | Reverse: 5′-AAAATTGATGGAATGCTAATTGGT-3′ |
| Gapdh | Forward: 5′-GGACTGGATAAGCAGGGCG-3′ |
|  | Reverse: 5′-TTTTGTCTACGGGACGAGGC-3′ |
| mTOR | Forward: 5′-GCCACTCTCTGACCCAGTTC-3′ |
|  | Reverse: 5′-ATGCCAAGACACAGTAGCGG-3′ |
| Phf8 | Forward: 5′-TGGGAGCATGCTTCAAGG-3′ |
|  | Reverse: 5′-GATTTCAAAGCAGGGTCATCA-3′ |
| mTOR upstream TSS* | Forward: 5′-TTGCCAACTGGTGCTCGTTT-3′ |
|  | Reverse: 5′-AAGAATTGGAGCTCGGGACC-3′ |
| mTOR TSS* | Forward: 5′-GGATGTTCCTCCCCAATCTTCG-3′ |
|  | Reverse: 5′-CAGACCCACCTAACTGACCGT-3′ |
| mTOR downstream TSS* | Forward: 5′-TAGGGGGCAGATCCCGAAAC-3′ |
|  | Reverse: 5′-CACTGTAGCTGTAACTCACAC-3′ |
| * TSS, transcriprtion start site | |
